# The Lifespan Trajectories of Brain Activities Related to Cognitive Control

**DOI:** 10.1101/2023.08.20.554018

**Authors:** Zhenghan Li, Isaac T. Petersen, Lingxiao Wang, Joaquim Radua, Guo-chun Yang, Xun Liu

## Abstract

Cognitive control plays a pivotal role in guiding human goal-directed behavior. While existing studies have documented an inverted U-shaped trajectory of cognitive control both behaviorally and anatomically, little is known about the corresponding changes in functional brain activation with age. To bridge this gap, we conducted a comprehensive meta-analysis of 129 neuroimaging studies using conflict tasks, encompassing 3,388 participants aged from 5 to 85 years old. We have three major findings: 1) The inverted U-shaped trajectory is the predominant pattern; 2) Cognitive control-related brain regions exhibit heterogeneous lifespan trajectories: the frontoparietal control network follows inverted U-shaped trajectories, peaking between 24 and 40 years, while the dorsal attention network demonstrates no clear trajectories; 3) Both the youth and the elderly show weaker brain activities and greater left laterality than young to middle-aged adults. These results reveal the lifespan trajectories of cognitive control, highlighting heterogeneous fluctuations in brain networks with age.

**Classification:** Biological Sciences/Psychological and Cognitive Sciences

## Introduction

The cognitive abilities of human beings dynamically change throughout the entire lifespan, experiencing rapid development in the early stages, and gradual decline in the later period of life. As one of the most fundamental cognitive functions, cognitive control is deeply engaged in various domains of high-level capabilities that humans greatly outperform other species, such as decision making, planning and problem solving^1^. Cognitive control refers to the cognitive processes that enable individuals to manage and regulate their attention, thoughts, and actions, which plays a vital role in goal-directed behavior, allowing us to focus on the target and ignore distractors^2,3^. For instance, cognitive control enables us to concentrate on reading in a library despite the presence of people chatting nearby.

Cognitive control provides fundamental support for normal human behaviors. Young adults typically maintain an optimal mature level of cognitive control^4^. However, the youth (including children and adolescents, less than 18 years) and elderly (60 years and older) individuals may struggle with behavioral problems because of their suboptimal cognitive control system^5–10^. While the state-of-art progress of cognitive/behavioral changes has been well-documented and shaped how the diagnosis of developmental/ageing related disorders^11^, the change of related neural system has been under-investigated. Understanding how the neural underpinning of cognitive control changes over the lifespan can yield valuable insights into the developmental and aging mechanisms of the human brain. This knowledge may assist in customizing cognitive training strategies based on related brain regions and their activities^12^.

Researchers generally believe that cognitive control ability follows an inverted U-shaped trajectory across the lifespan^5,13,14,17^. This inverted U-shaped trajectory has been generally supported by behavioral and anatomical evidence. The Eriksen Flanker task (requiring participants to respond to central stimuli while ignoring flanking distractions) has been widely utilized to detect cognitive control across the lifespan^15^, and results suggest a clear U-shape trajectory of conflict cost (measured by worsened behavioral performance, e.g., reaction time, in incongruent compared to congruent conditions) with age^16–18^. Similar results have been observed in other conflict tasks, such as color-word Stroop (requiring participants to name the ink color of a word that is incongruent with the word’s semantic meaning)^16^. Recent large-cohort studies have also found that gray and white matter volumes across all brain regions exhibit overall inverted U-shaped trajectories with age, with the gray matter volume peaking at early adolescence and the white matter volume peaking at young adulthood^19,20^. During late adulthood, normal ageing yields a protracted decline of brain structure, with the volumes of both gray matter and white matter reduced^14^. Consistently, the gray matter volumes of the frontal and parietal regions, which are essential in cognitive control tasks^21^, have also been found to increase during early childhood and atrophy in early elderly age^22^.

However, it remains largely unknown how brain activities related to cognitive control change over the lifespan. Previous research has primarily focused on brain activities in either youths or elderly adults, rather than examining changes across the entire lifespan. With conflict paradigms, children and adolescents are often found to have lower brain activity than young adults in frontoparietal regions (refs.^23–28^, but see Bunge et al. ^29^). For example, a study utilizing the Flanker task revealed that children aged 8−12 years had reduced activation in dorsolateral prefrontal regions compared to young adults, suggesting an immature cognitive control system^27^. However, brain activity differences in elderly adults as compared to young adults during cognitive control tasks have been less consistently reported^14^. Some studies have found that elderly adults have lower neural activity in frontoparietal regions than young adults^30–32^, possibly because elderly adults may be unable to engage in an equal level of control-related activity due to functional decline. On the other hand, other studies have found that elderly adults may exhibit greater brain activity in frontoparietal regions than young adults^33,34^, possibly because they have recruited additional brain regions to compensate for their decreased efficiency in utilizing control resources. Adding to the debate, it has been proposed that the cognitive control function in elderly adults might not decline at all^35,36^. These conflicting findings underscore the complexity of understanding age-related differences in cognitive control.

Few studies have directly tested the change of brain activities related to cognitive control across the lifespan. One existing study observed a positive association between the activation of the bilateral prefrontal cortex and age^37^. Given the relatively small sample size (N = 30), the reliability of these findings is somewhat limited. As a result, it is difficult to draw a clear conclusion about how brain activities related to cognitive control change across the entire age range.

One direct way to test the lifespan trajectory of brain activation related to cognitive control is to conduct a large cohort of neuroimaging study with participants covering a wide age range. To the best of our knowledge, such studies have not yet been conducted. An alternative approach is to utilize meta-analyses to combine the results of existing studies targeting different age groups. Compared to the large cohort studies, meta-analyses are more accessible and resource-saving. In addition, meta-analyses can increase the statistical power and generalizability by combining various studies, reducing heterogeneity and bias from individual studies’ methods, populations, or confounding variables^38^. Importantly, meta-analyses can reveal patterns not evident in individual studies, like nonlinear effects or interactions^38^. Several neuroimaging meta-analyses have examined age-related changes of cognitive control brain activity^31,39,40^, but limitations such as incomplete age coverage and insufficient studies in certain age ranges prevent them from appropriately answering questions about the lifespan trajectory of cognitive control functions^31,39,40^. These studies have primarily focused on spatial convergence and/or diversity of coordinates across different age ranges, offering limited insights into lifespan trajectories of activity strength. Although a few studies have attempted parametric meta-regression to examine the age-related differences in cognitive control-related brain activity^39,41^, they have been constrained by utilizing linear models that may overlook non-linear trajectory patterns, such as the inverted U-shaped trend.

The goal of this study is to provide a comprehensive examination to reveal the lifespan trajectory of brain activities responsible for cognitive control. Instead of encompassing various aspects of cognitive control, we focus on conflict processing for several reasons. First, conflict processing reflects the fundamental cognitive control ability to maintain a goal while avoiding distractions^2^. Second, its mechanisms in young adults are relatively well-known, with the frontoparietal and cingulo-opercular networks engaged^21,42^, providing a baseline reference for our study. Third, conflict tasks with neuroimaging data have been widely applied to both younger and elderly groups, making a systematic meta-analysis feasible. Lastly, different conflict tasks share key components of cognitive control, such as conflict monitoring^43^ and inhibitory control^8^, which enables us to conduct effect-size-based meta-analyses using the congruency effect (i.e., the contrast between incongruent and congruent/neutral conditions). Including other sub-processes of cognitive control may introduce heterogeneity and make effect sizes incomparable.

Previous research suggests that better cognitive control-related performance is often associated with greater brain activity, especially in the prefrontal cortex^44^. Therefore, we hypothesized that brain activities related to cognitive control might follow an inverted U-shaped trajectory like that of behavior patterns. Additionally, we hypothesized that different brain networks may show some heterogeneity in their lifespan trajectories.

## Results

### Sample Description

A total of 3,611 articles were identified including 3,484 articles searched from the database, 111 articles adopted from a previous study^21^, and 16 articles searched from the references of crucial articles. After excluding duplicates and applying exclusion criteria, 119 articles including 129 studies with 3,388 participants and 1,579 brain activation foci, were included in this meta-analytic study (Supplementary Fig. S1 and Table S1). The average age of participants ranged from 8 to 74 years, with the individual age ranging from 5 to 85 years. A demonstration of the age distribution for each included study is presented in Supplementary Fig. S2.

### Regions Related to Cognitive Control Identified by both ALE and SDM

To enhance the replicability and robustness of our finding and enable direct comparison with a previous study^21^, we conducted the mean analysis across all studies with the activation likelihood estimation (ALE) and seed-based d mapping (SDM) approaches separately. First, we performed the single dataset analysis with the GingerALE software to explore the brain regions consistently reported in all the included studies (see section “*Activation Likelihood Estimation (ALE)*” in Methods). Results showed significant activation in the frontoparietal regions, including the left dorsolateral prefrontal cortex, bilateral frontal eye field, right inferior frontal gyrus and bilateral inferior parietal lobule; the cingulo-opercular regions, including the supplementary motor area and bilateral insula; and other regions, including the left inferior temporal gyri (Fig. 1A and Supplementary Table S2). Second, we calculated the average brain activation based on the effect sizes reported in all studies using the SDM with permutation of subject images (SDM-PSI) software (see section “*Mean Analyses Across all Studies*” in Methods). Similar to the ALE results, we found significant activation in the frontoparietal regions (left inferior parietal lobule, right inferior frontal gyrus, and right middle frontal gyrus), the cingulo-opercular regions (left anterior cingulate cortex), and other regions including bilateral inferior temporal gyrus, right caudate nucleus, right cerebellum, and left anterior thalamic projections (Fig. 1B and Supplementary Table S3). The count of voxels revealed that the overlapped area (3,496 voxels, Fig. 1C) accounted for 96.3% of the regions from the ALE analysis (3,635 voxels), and 15.0% of the regions from the SDM analysis (23,240 voxels). This result suggested that SDM analysis could be a reliable approach for detecting brain regions, thereby laying a solid foundation for using this approach in subsequent analyses. In addition, a robustness analysis suggested that age does not influence the mean results of SDM analysis (Supplementary Fig. S3 and Table S2).

**Fig. 1.**
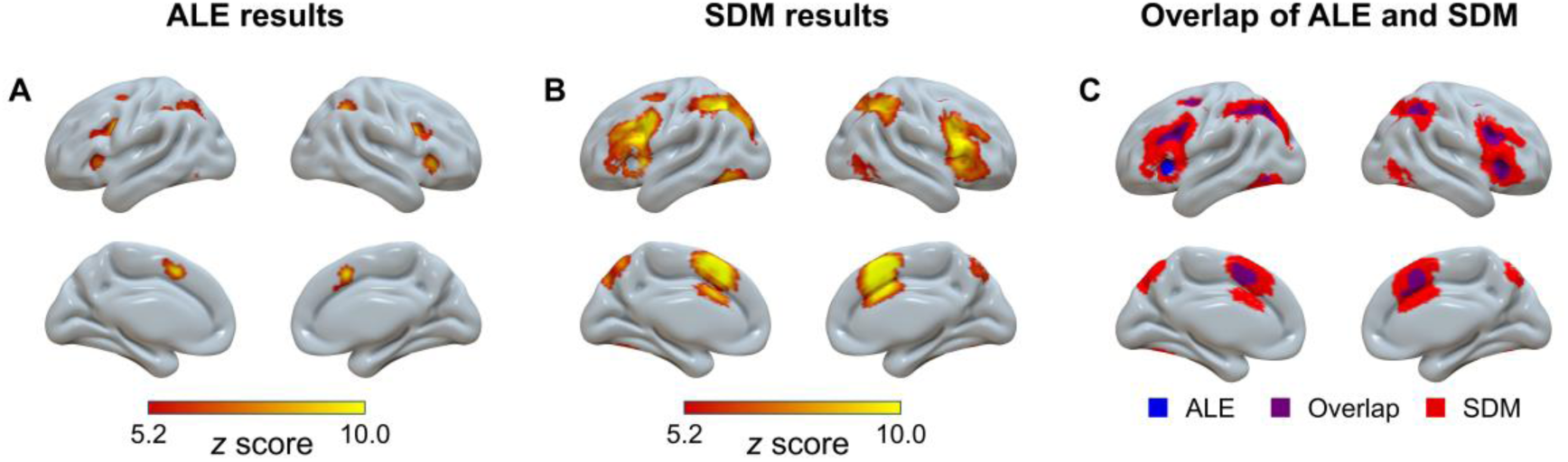
Overview of significant clusters across all studies regardless of age in the ALE meta-analysis (A), the SDM meta-analysis (B), and their overlap (C). ALE = activation likelihood estimation; SDM = seed-based *d* mapping.

### Trajectories of Cognitive Control Regions Identified in the Mean Analysis

Having identified nine brain regions in the mean analysis using SDM-PSI, we proceeded to explore how the activation levels of these regions change with age. To this end, we extracted brain activity data from all studies for each identified region. Before performing the meta-regression, we verified the effectiveness of using mean age as a predictor with a simulation approach (Supplementary Note S1 and Fig. S4). We then subjected the extracted data to separate generalized additive model (GAM) analyses, factoring in the confounding covariates (see section “*Generalized Additive Model (GAM) Fitting*” in Methods). The analyses (Supplementary Table S4) revealed significant age-related changes in the activation levels of 4 out of the 9 regions (Fig. 2, l-ACC, r-IFG, l-ITG, and r-CN). Visualization of the trajectories suggests that these regions showed inverted U-shaped patterns. The significance of their inverted U-shaped trajectories was further examined using a two-line test approach^45^. Results suggest that all four regions involve an increase in activity from childhood to young adulthood (approximately up to 30 years of age), and three of them (r-IFG, l-ITG and r-CN) showed consistent decrease in the later stages of age (Supplementary Note S2). The other five regions (Fig. 2, l-IPL, r-ITG, r-MFG, r-CB and l-ATP) showed no significant age-related changes. Notably, none of the clusters showed significant publication bias based on Egger’s test (*p*s > 0.86), and they all showed low between-study heterogeneity (*τ*s < 0.13, *Q*s < 12.11, *I*²s < 25%). This indicates that the observed results are not likely influenced by biased reporting or substantial variability in the included studies. A robustness analysis further ruled out the potential influence of including unpublished and non-English studies in this study (Supplementary Note S4 and Fig. S5).

**Fig. 2.**
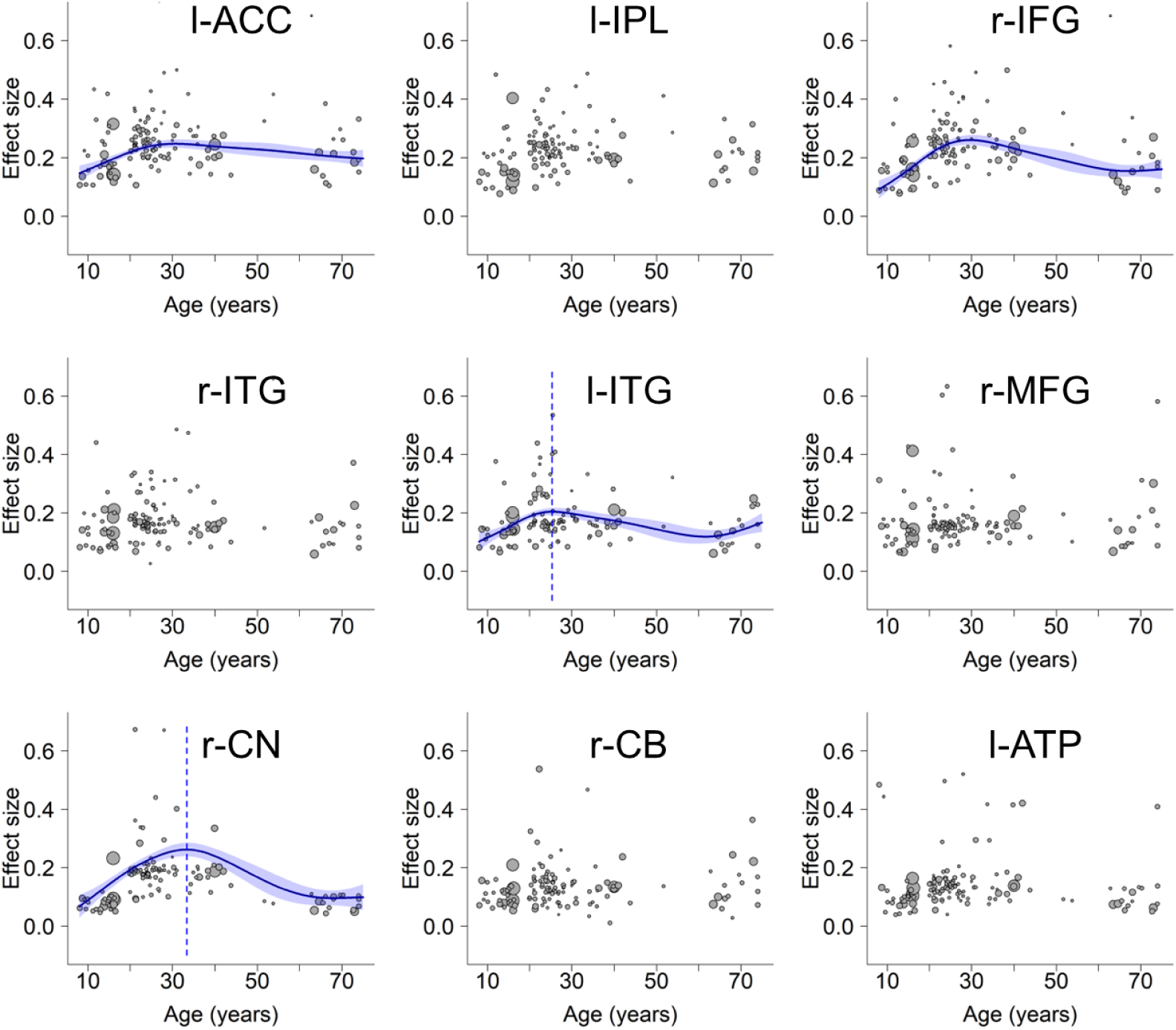
Lifespan trajectories within regions identified in the mean analysis. l-ACC: left anterior cingulate cortex, l-IPL: left inferior parietal lobule, r-IFG: right inferior frontal gyrus, r-ITG: right inferior temporal gyrus, l-ITG: left inferior temporal gyrus, r-MFG: right middle frontal gyrus, r-CN: right caudate nucleus, r-CB: right cerebellum, l-ATP: left anterior thalamic projections. Scattered plots are the effect sizes as a function of age, with curves fitted by GAM. The sizes of the scattered dots show the square root of model weights (1/variance) for each study. Shaded areas around the curves represent standard errors. Dashed lines indicate peak ages. Panels l-ACC and l-IPL do not show the peak age due to an insignificant decrease at the later stage (Supplementary Note S2).

Note that some regions are large, containing up to over 9,000 voxels. This could obscure the distinct trajectories specific to different subregions, which are crucial in understanding the hierarchical patterns of lifespan trajectories. We address this by using a more granular approach below.

### Detecting Different Trajectories of Whole Brain Activities

To explore various possibilities of lifespan trajectories, we grouped studies by their mean age into the youth, young to middle-aged, and elderly groups, and then conducted several contrast analyses (see section “*Contrast Analysis*” in Methods). These analyses did not reveal any regions that exhibited significantly higher or lower brain activity in the youth compared to others (the combination of young to middle-aged and older adults) (Table 1). Similarly, we did not observe any regions with higher or lower brain activity in older adults compared to others (the combination of the youth and young to middle-aged adults) (Table 1). These results excluded the possibilities of increase/decrease-then-stable and stable-then-increase/decrease trajectories (Fig. 3, panels D, F, G, and I). In addition, we failed to observe any region showing lower activity in young to middle-aged adults than others (the combination of the youth and older adults), and thereby ruled out the possibility of the upright U-shaped trajectory (Table 1, Fig. 3E).

**Table 1.**
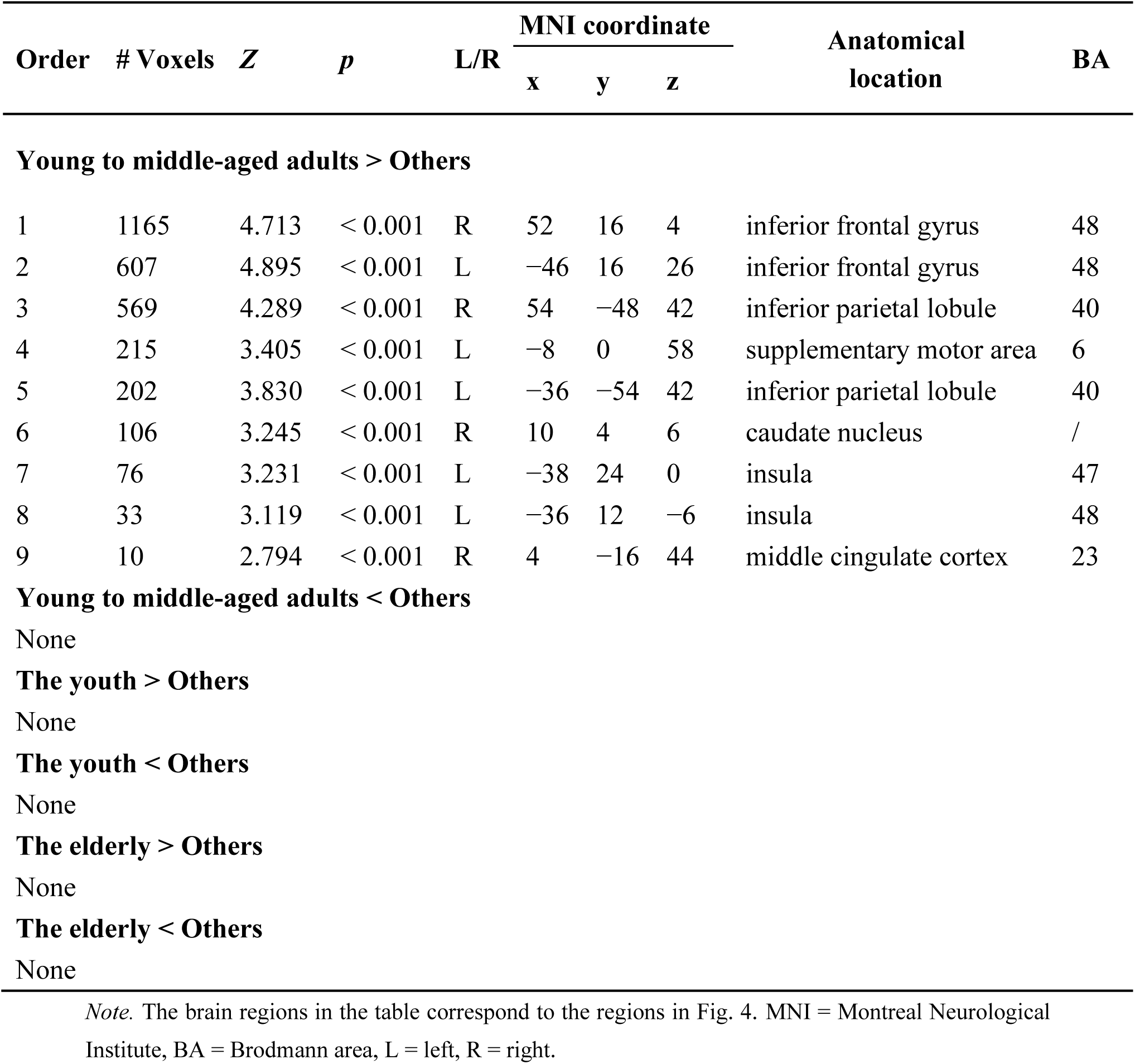
Brain areas activated in the contrast of one age group versus others (voxelwise FWE-corrected, *p* < 0.001, with minimum cluster size ≥ 10 voxels) with the SDM-PSI.

| Order | # Voxels | Z | p | L/R | MNI coordinate |  |  | Anatomical location | BA |
| --- | --- | --- | --- | --- | --- | --- | --- | --- | --- |
|  |  |  |  |  | x | y | z |  |  |
| Young to middle-aged adults > Others |  |  |  |  |  |  |  |  |  |
| 1 | 1165 | 4.713 | < 0.001 | R | 52 | 16 | 4 | inferior frontal gyrus | 48 |
| 2 | 607 | 4.895 | < 0.001 | L | −46 | 16 | 26 | inferior frontal gyrus | 48 |
| 3 | 569 | 4.289 | < 0.001 | R | 54 | −48 | 42 | inferior parietal lobule | 40 |
| 4 | 215 | 3.405 | < 0.001 | L | −8 | 0 | 58 | supplementary motor area | 6 |
| 5 | 202 | 3.830 | < 0.001 | L | −36 | −54 | 42 | inferior parietal lobule | 40 |
| 6 | 106 | 3.245 | < 0.001 | R | 10 | 4 | 6 | caudate nucleus | / |
| 7 | 76 | 3.231 | < 0.001 | L | −38 | 24 | 0 | insula | 47 |
| 8 | 33 | 3.119 | < 0.001 | L | −36 | 12 | −6 | insula | 48 |
| 9 | 10 | 2.794 | < 0.001 | R | 4 | −16 | 44 | middle cingulate cortex | 23 |
| Young to middle-aged adults < Others |  |  |  |  |  |  |  |  |  |
| None |  |  |  |  |  |  |  |  |  |
| The youth > Others |  |  |  |  |  |  |  |  |  |
| None |  |  |  |  |  |  |  |  |  |
| The youth < Others |  |  |  |  |  |  |  |  |  |
| None |  |  |  |  |  |  |  |  |  |
| The elderly > Others |  |  |  |  |  |  |  |  |  |
| None |  |  |  |  |  |  |  |  |  |
| The elderly < Others |  |  |  |  |  |  |  |  |  |
| None |  |  |  |  |  |  |  |  |  |

*Note.* The brain regions in the table correspond to the regions in Fig. 4. MNI = Montreal Neurological Institute, BA = Brodmann area, L = left, R = right.

**Fig. 3.**
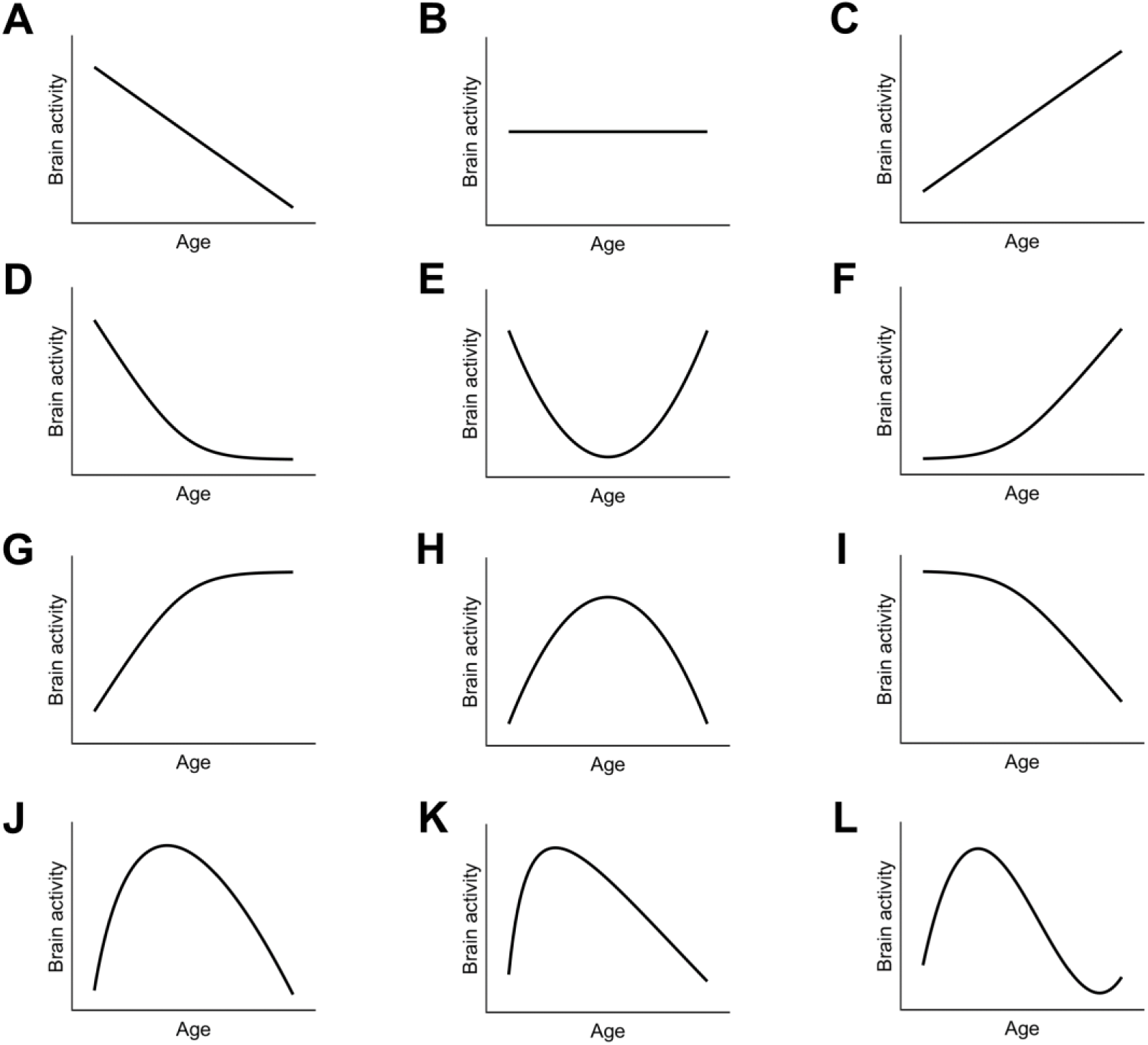
The lifespan trajectories explored in our study. Panels A-C show linear decrease, flat, and linear increase patterns, respectively, and were modelled with the linear function. Panels E and H show the upright and inverted U-shapes, respectively, and were tested with the contrast between young to middle-aged adults and others, as well as with the quadratic function. Panels D, F, G, and I show combinations of a stable period and an increase/decrease period across the lifespan, and were tested with the contrast between the youth and others, or between the elderly and others. Panels J, K and L show the variants of inverted U-shaped trajectories, which capture the possibly early peak feature. They were tested with square root, quadratic logarithmic, and cubic functions, respectively. See Methods for detailed models.

However, we identified greater activity in young to middle-aged adults compared to others in the frontoparietal regions, including bilateral inferior frontal gyrus and bilateral inferior parietal lobule; the cingulo-opercular regions, including left supplementary motor area, left insula, and right middle cingulate cortex; and a subcortical region—right caudate nucleus (Fig. 4 and Table 1). This result essentially supports an inverted U-shaped trajectory. Notably, none of the clusters showed significant publication bias based on Egger’s test (*p*s > 0.79), and they all showed low between-study heterogeneity (*τ*s < 0.17, *Q*s < 12.21, *I*²s < 25%). This indicates that the observed results are not likely influenced by biased reporting or substantial variability in the included studies. Consistently, further contrast analyses revealed that the young to middle-aged adults showed greater activity than both the youth (Supplementary Fig. S6) and the elderly (Supplementary Fig. S7).

**Fig. 4.**
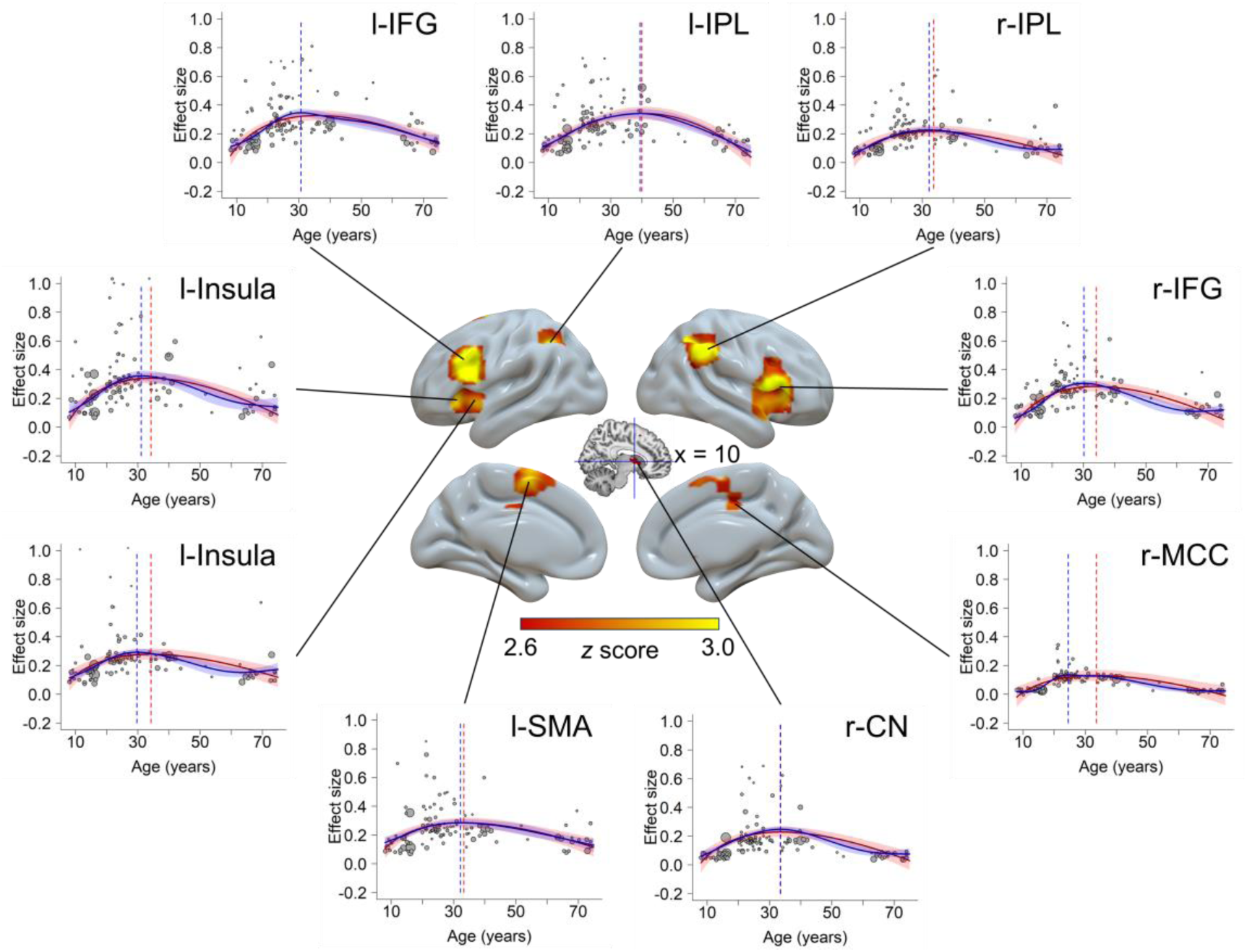
Brain regions showing inverted U-shaped trajectory patterns. Scattered plots are the effect size as a function of age, with curves fitted by GAM (blue color) and the best simplified model (red color). Shaded areas around the curves represent standard errors. Dashed vertical lines show peak ages estimated from GAM (blue) and simplified model (red). The sizes of the scattered dots show the square root of model weights (1/variance) for each study. r-IFG: right inferior frontal gyrus, l-IFG: left inferior frontal gyrus, r-IPL: right inferior parietal lobule, l-SMA: left supplementary motor area, l-IPL: left inferior parietal lobule, r-CN: right caudate nucleus, l-Insula: left insula, r-MCC: right middle cingulate cortex.

### Fitting the Lifespan Trajectories with the GAM

For each region identified from the contrast between young to middle-aged adults and others, the GAM could fit the data significantly with a smooth curve (Fig. 4), with degrees of freedom varying from 2.9 to 6.7. Peak ages of the inverted U-shaped trajectories were between 24.5 and 39.4 years. Detailed statistics are shown in Supplementary Table S4.

### Model Simplification of the Lifespan Trajectories

Considering the GAM may overfit the data, we fitted the results with simpler models commonly adopted in lifespan developmental literature^46–48^, including the quadratic, cubic, square root and quadratic logarithmic models, and estimated the goodness of fit by comparing their Akaike information criterion (AIC) (see section “*Model Simplification and Model Comparison*” in Methods). Results showed that the quadratic model provided the best fit for capturing the age-related changes in left inferior parietal lobule, while the square root model demonstrated the best goodness of fit for all the other regions (Supplementary Table S5). We also calculated the peak age for each region based on the optimal model, and results showed that the peak ages ranged from 33.3 to 40.0 years. In addition, we found the peak ages obtained from the above optimal model (i.e., square root or quadratic models) and the GAM are consistent, *r* = 0.71, *p* = 0.032, 95% CI = [0.09 0.93]. See Fig. 4 and Supplementary Table S6 for details.

### Fitting the Whole Brain with Square Root and Linear Models

Based on model comparisons, we found that the square root model provided the best goodness of fit for the age-related change of the brain activities. To supplement the inverted U-shaped results from the contrast analysis, the square root function was then submitted to whole-brain meta-regression analyses in SDM-PSI (see section “*Meta-regression Analyses*” in Methods).

By fitting the activation over the whole brain, we found seven significant brain regions, including the bilateral inferior frontal gyrus, right angular, left inferior parietal lobule, right insula, right caudate nucleus, and left anterior thalamic projections (Fig. 5A and Supplementary Table S7). These results further supported the existence of the inverted U-shaped regions we initially identified (Fig. 4).

**Fig. 5.**
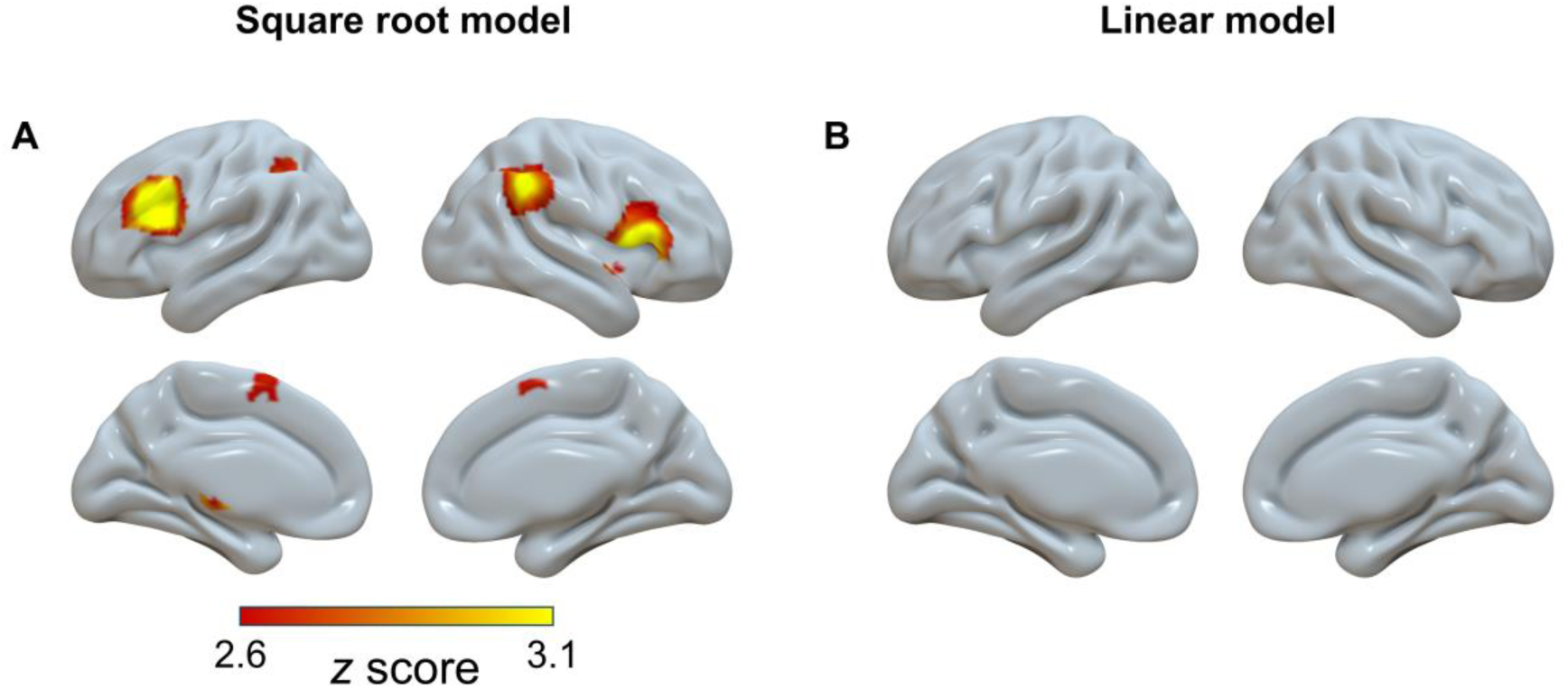
Significant clusters (voxel-wise FWE-corrected, *p* < 0.001, voxels ≥ 10) showing square root pattern (A) and linear pattern (B) with age in the model fitting.

In addition, we explored the whole-brain trajectories with a linear meta-regression (see section “*Meta-regression Analyses*” in Methods). However, no significant regions were observed (Fig. 5B), even under a more tolerant threshold of uncorrected *p* < 0.01.

### Dissociated Brain Networks with Distinct Lifespan Trajectories

We note that the inverted U-shaped regions constitute only part of the cognitive control-related regions (Fig. 2), and the remaining regions show a less clear trajectory. To further elucidate the spatial distribution of brain regions following these different trajectory patterns, we used the results from the mean SDM analysis (Fig. 1B) as the mask and replotted the results of the contrast analysis between young to middle-aged adults and others with two different thresholds, and then compared them with the Yeo’s 7-network atlas^49^ (Fig. 6). Visualization of the spatial distribution patterns revealed a dissociation between the middle frontal gyrus and its adjacent rostral and caudal areas (Fig. 6A). Moreover, the distribution of inverted U-shaped regions was more consistent with the frontoparietal control network (FPCN), and the distribution of non-U-shaped regions was more closely related to the dorsal attention network (DAN, Fig. 6B). The count of voxels revealed that a numerically larger portion of the inverted U-shaped regions overlapped with the FPCN (2,538 voxels) than with the DAN (932 voxels), while the overlap with the cingulo-opercular network (CON) was in between (1,867 voxels). Conversely, a numerically larger portion of the non-U-shaped regions overlapped with the DAN (3,009 voxels) than with the FPN (2,447 voxels), while the overlap with the CON was lower (1,594 voxels).

**Fig. 6.**
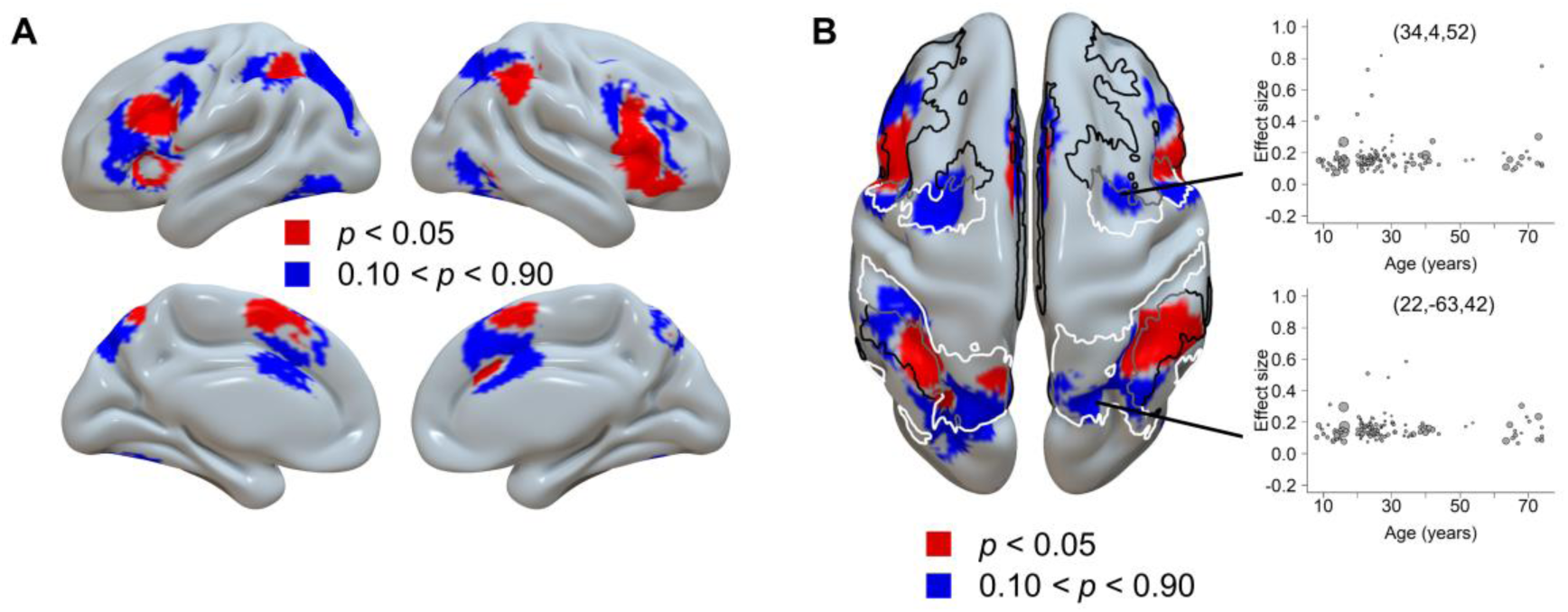
Dissociated brain regions based on their trajectory patterns. A) The regions following inverted U-shaped trajectories (red color) and non-U-shaped trajectories (blue color). B) The axial view of the same results. The border lines display the frontoparietal control network (black), dorsal attention network (white), and their boundary (gray) from Yeo’s 7-network atlas^49^. Cingulo-opercular network was not plotted due to its less clear dissociation among the two maps. The two scatter plots show two example regions showing the non-U-shaped trajectory, one ([34, 4, 52]) representing a peak region from the average brain activity analysis (Supplementary Table S3), and the other ([22, −63, 42]) representing a region displaying a weak age-related change from the contrast analysis with p between 0.49 and 0.51. The GAM analysis showed that neither coordinate could be adequately fitted by a smoothed curve, with *p*s > 0.22.

To further investigate the potential functional difference between the two sets of brain regions, we decoded the related terms with the Neurosynth decoder^41^ (see section “*Neurosynth Decoding Analysis*” in Methods). Results showed that the inverted U-shaped regions were related to the term “cognitive control”(*r* = 0.187), but were less so to “attentional” (*r* = 0.100) and “monitoring” (*r* = 0.072). Note the keyword “monitoring” refers to the major function of the cingulo-opercular network^21^. On the other hand, the non-U-shaped regions were related to the term “attentional” (*r* = 0.240) but were less so to “monitoring” (*r* = 0.090) and “control” (*r* = 0.062) (Supplementary Table S8). This result suggests that the inverted U-shaped and non-U-shaped regions may be associated with cognitive control and attention, respectively, which is consistent with the frontoparietal control network and dorsal attention network as identified in the atlas overlapping analysis.

### Lifespan Trajectory of the Laterality

We also tested how the laterality of the brain activity changes with age (see section “*Laterality Analysis*” in Methods). We first modeled the laterality trajectory using the GAM. Results showed a significant model fitting, *F*(3.0, 3.7) = 3.49, *p* = 0.012, *R*^2^ = 0.24. Moreover, we fitted the data with the four simplified models (i.e., the quadratic, cubic, square root, and quadratic logarithmic models). The results showed that a square root function provided the best goodness of fit, with the *β*_sqrt(age)_ = −0.68 (95% CI = [−1.02, −0.33]), *p* < 0.001. The two-line test suggests the hypothetical peaks from both models did not reach significance (Supplementary Note S2). A visually upright U-shaped trajectory indicated the youth and elderly adults tended to be more left-lateralized across the whole brain. A further comparison of the relative levels of laterality across different age groups revealed that both the youth and the elderly groups exhibited greater left lateralization than the young to middle-aged adult group (Supplementary Note S3). This left-lateralized pattern could be illustrated by the brain map estimated with voxel-wise laterality calculation (Fig. 7).

**Fig. 7.**
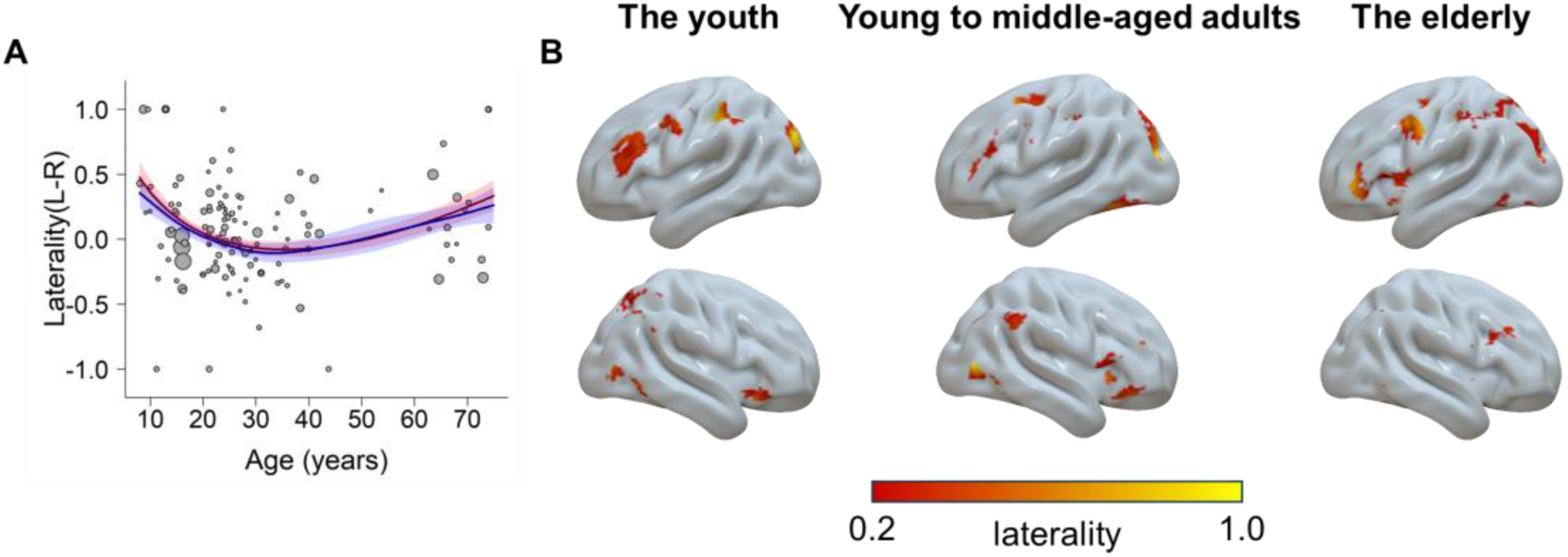
The laterality as a function of age. A) The trajectory fitted with a square root model (red) and the GAM (blue). Higher values mean more left-lateralized and lower values mean more right-lateralized. B) Visualization of the laterality for each group. Regions in the left hemisphere show the left laterality, and vice versa.

## Discussion

The present study yielded three primary findings: 1) Among different possible trajectories, only the inverted U-shaped trajectories were reliably observed across the whole brain; 2) The cognitive control-related brain regions exhibit heterogeneous lifespan trajectories: the frontoparietal control network (such as the inferior frontal gyrus and inferior parietal lobule) follows inverted U-shaped trajectories, peaking between 24 and 40 years, while the dorsal attention network (such as the frontal eye field and superior parietal lobule) demonstrates less clear trajectories with age; 3) The youth and the elderly demonstrate weaker brain activities and a relatively greater extent of left laterality compared to the young to middle-aged adults. These results provide strong evidence for the existence of cognitive control regions exhibiting inverted U-shaped trajectory, and also show the heterogenous lifespan trajectories in different brain regions.

### The Inverted U-shaped Trajectory of Brain Activity Related to Cognitive Control

The main finding is that a wide range of cognitive control regions follow inverted U-shaped lifespan trajectories, but no regions showed decrease-then-stable (Fig. 3D), up-right U-shaped (Fig. 3E), stable-then-increase (Fig. 3F), increase-then-stable (Fig. 3G), stable-then-decrease (Fig. 3I), or linear trajectories (Fig. 3A and 3C).

The greater activation in the frontoparietal control network among young to middle-aged adults compared to the youth and the elderly supports the notion that cognitive control abilities may not be fully developed in children and may decline in the elderly. This finding is consistent with the idea that the cognitive control system is most effective in young adulthood, suggesting a possible correlation between the higher functional activations in the brain and the superior performance of young adults on cognitive control tasks^50^. Consistently, a previous study^29^ showed that behavioral performance (success of interference suppression) is positively correlated with the activity in frontal regions. Similar patterns have been repeatedly reported^51,52^, although the opposite results have also been observed^53^.

The inverted U-shaped trajectory of brain activation might be associated with the development of brain structure and functional changes. First, it may reflect the well-documented structural changes that occur in these regions across the lifespan, which include synaptic pruning, myelination, cortical thinning, and white matter maturation^19,54^. For example, the density of dopamine receptors increases during adolescence and young adulthood and subsequently declines with age^55^. These changes can affect the efficiency and connectivity of neural circuits within the frontoparietal control network^13,56^. Second, the inverted U-shaped trajectory may also arise from functional changes resulting from the modulation of neurotransmitters, hormones, and environmental factors^57^. Understanding change patterns of brain structure and function is critical for developing interventions and treatments aimed at improving cognitive control abilities across the lifespan.

The present results further revealed that the inverted U-shaped lifespan trajectories of cognitive control regions are not uniform. The GAM fitting results (Fig. 4, Supplementary Table S4) showed that the subregions exhibited varying degrees of association with age. Moreover, the model simplification demonstrated different underlying trajectory curves, with most regions showing a skewed shape that could best be fitted with a square root model, except that one region showed a symmetric quadratic shape. The quadratic lifespan trajectory has been well-documented in previous studies^19,46,48,58^, while the application of the square root model has been relatively rare^59^. The square root model can better capture the early peak in the trajectory. We also identified different peak ages for those regions, ranging from 24 to 40 years, suggesting that cognitive control regions may not develop at the same rate.

### Hierarchical development trajectories in different brain networks

Previous research has indicated that the attentional orientation function is preserved during ageing^35,60^. Consistently, we found that the dorsal attention network regions underlying the attentional orientation showed no significant age-related change, in contrast to the inverted U-shaped trajectory in frontoparietal control network regions. In addition, we observed that the supporting regions mediating top-down control with motor^61^ (right cerebellum, Fig. 2) and sensory^62^ (left anterior thalamic projections, Fig. 2) functions also lack the sensitivity to age. The dissociation across regions may reflect hierarchical associations with age on brain function.

The brain regions are organized in a functional hierarchy, with the frontoparietal control network at the highest level. It acts as a hub that interacts with other systems, including the dorsal attention network^63^. The cingulo-opercular network did not present a clear dissociation of trajectory patterns, possibly suggesting its intermediate position between the frontoparietal control and dorsal attention networks^64^. During conflict tasks, these networks function in a hierarchical manner. The frontoparietal control network maintains task goals and resolves conflicts, the cingulo-opercular network monitors conflict, and the dorsal attention network directs attention towards task-relevant stimuli^65^. Even within the prefrontal cortex itself, a hierarchical organization exists, with middle frontal areas occupying the peak position^66^. This is in line with our finding that the middle frontal cortex is dissociated from rostral and caudal frontal regions (Fig. 6A). In addition, previous research suggests that the frontoparietal control network can be further divided^67^, with the rostral and caudal frontal regions observed in our study aligning closely with the sub-network that connects more strongly with the dorsolateral attention network. This may explain why some areas within the frontal region do not show age-related changes.

Furthermore, different brain regions exhibit different age-related changes. Higherorder regions typically have more complex lifespan trajectories^58^ and reach peaks during later periods^56,68^. Specifically, prefrontal control regions are among the last to mature and one of the earliest to decline^5,13,69^. Therefore, the different lifespan trajectory patterns among different networks likely reflect their hierarchical positions of age-related changes.

### Implications for the Compensatory and Asymmetry Reduction Theories

Critically, there was no region showing higher activity in the elderly compared to the young to middle-aged adults, but we observed several regions showing the opposite (Supplementary Note S3 and Table S9). The results persisted after we controlled the behavioral congruency effect (Supplementary Note S4 and Fig. S8), thereby ruling out the possibility of weaker brain activity associated with poor behavioral performance in the elderly. The observed decrease in brain activation among the elderly might be attributed to several interrelated factors. First, cognitive control regions, especially the frontal area, tend to shrink with age, leading to a reduction in overall brain volume and potential loss of synaptic integrity^70^. This shrinkage can impair the brain’s ability to effectively process and manage complex tasks. Another significant factor is the impairment of neurovascular coupling, the relationship between neuronal activity, synaptic function, and subsequent blood flow, which disrupts the brain’s ability to maintain optimal function during cognitive tasks^71,72^. Furthermore, the decrease in cerebral blood flow with age can diminish the delivery of essential nutrients and oxygen to the brain, impairing its overall functionality^73^. These changes could lead to regional abnormalities, such as blood flow, blood volume, metabolic rate, or BOLD-derived physiologic proxies like the fractional amplitude of low-frequency fluctuation and regional homogeneity^74^. Future studies may validate and extend our study by adopting the age-sensitive regions we observed and testing other measurements, such as resting-state data, which are more easily collected in large-scale studies involving children and the elderly compared to task-based activations.

The compensatory theory^10^ proposes that the elderly recruit additional brain regions to compensate for age-related cognitive decline, but our results did not show this pattern. We suggest that the absence of compensatory upregulation in frontoparietal regions among the elderly observed in our study might be attributed to limited available resources when cognitive control related brain regions are already fully engaged^75,76^. Previous research has shown that younger adults recruit lower activity in frontoparietal regions during the congruent condition but significantly greater activities during the incongruent condition. In contrast, older adults already show a relatively higher activation during the congruent condition, leaving limited capacity for further increases in activation during the incongruent condition^32^. This is consistent with our findings, which are based on the contrast between incongruent and congruent conditions. Moreover, the nature of the task investigated might influence whether there is an upregulation in cognitive control regions with age. Upregulation in the frontal regions usually compensates for memory and sensory declination due to deficits in the hippocampus and sensory cortices^77^. Semantic cognition^78^ might also be a target of compensation. However, conflict tasks seem to rely minimally on memory, and involve relatively simple sensory stimuli (e.g., colors and locations) and simple semantic processing (e.g., reading a word). As such, conflict tasks may not necessitate compensation in these functions.

In addition, compensation in older adults may manifest as increased recruitment of bilateral regions and homologues with age^79^. For example, the hemispheric asymmetry reduction in older adults (HAROLD) theory^80^ suggests that older adults typically exhibit less lateralization, either as a compensatory response to functional deficits or as a reflection of neural dedifferentiation. However, we observed that the elderly showed greater left lateralization compared to young to middle-aged adults. This finding is inconsistent with the assumption of HAROLD but aligns with the right hemi-ageing model^81^, which posits that the right hemisphere is more vulnerable to age-related decline. Prior research has shown that functional connectivity within the frontoparietal control network is more disrupted in the right hemisphere than in the left during ageing^82^. This suggests that neural resources in the right hemisphere might be more limited for the elderly, reducing its capacity to compensate for cognitive demands. Stronger patterns of left laterality were also identified in childhood in the current study, primarily noticeable within the prefrontal region (Fig. 7), which may reflect the earlier development of the left hemisphere compared to the right^83^. In contrast, we found lower lateralization in young adults. It is possible that previous studies showing stronger laterality in young adults may have been biased by too small sample sizes and the use of non-quantitative methods for calculation of laterality^84,85^. Moreover, because both left and right lateralized results were reported in the literature on laterality^81^, it is reasonable to observe low laterality for young to middle-aged adults in the current meta-analysis.

### Methodology Implications

By incorporating all the studies, our results demonstrate that the SDM can reliably identify brain regions as the ALE. However, the SDM has the added advantage of fully utilizing existing effect size data and coordinates^86^, allowing us to compare the relative activity strength among various age groups, such as the contrast between young to middle-aged adults and elderly groups. More importantly, this approach allows for metaregressions to examine parametric relationships between brain region activity and age, providing insights into the lifespan trajectories of cognitive control regions.

### Limitation of Results

One caveat to consider in this study is the non-uniform distribution of age among the included studies. Specifically, there is a noticeable gap in the age range of 45 to 60 years. Consequently, the observed age distribution could potentially influence the results of the regression analysis. We hope that future research could allocate more attention to the middle-aged period, considering the significant cognitive and neural changes during this stage, such as the onset of cognitive decline^15,17,87^. In addition, it is crucial to avoid the occurrence of ecological fallacy^88^ (associations observed at the group level are erroneously assumed to apply to individuals) when interpreting the results of metaregression analyses. Therefore, associations between brain activities and age across various studies do not provide direct insights into the specific age-related changes at the individual level. Future research incorporating individual-level investigations (e.g., longitudinal follow-up studies) is crucial to obtaining a more comprehensive understanding of these relationships.

## Conclusions

Our meta-analysis adopted advanced meta-regression approaches to chart the lifespan trajectories of cognitive control brain activities. We observed inverted U-shaped changing patterns in regions aligned with the frontoparietal control network, with the peaks occurring between 24 and 40 years. In contrast, the dorsal attention network does not present a clear age-related trajectory. This dissociation may reflect the hierarchy of brain development in different regions. No other trajectory patterns were observed, highlighting the predominance of the inverted U-shaped pattern in the lifespan trajectory of cognitive control. Furthermore, we found the youth and elderly showed a more asymmetric brain distribution than young to middle-aged adults. In sum, these results demonstrate the multifaceted nature of age-related changes in cognitive control brain function.

## Methods

### Literature Preparation

#### Literature Search

We report how we determined all data exclusions (if any), all manipulations, and all measures in the study. We first searched both English and Chinese articles on the youth and the elderly from PUBMED, Web of Science and CNKI (China National Knowledge Infrastructure) till 2022. The following search terms were applied in titles, abstracts, table of contents, indexing, and key concepts: *(“Stroop” OR “Flanker” OR “Simon” OR “SNARC” OR “Navon” OR “interference” OR “cognitive conflict”) AND (“fMRI” OR “functional resonance imaging” OR “functional imaging” OR “neuroimaging” OR “PET”) AND (“children” OR “kids” OR “adolescents” OR “teenagers” OR “underage” OR “aged” OR “old” OR “older” OR “elder” OR “elderly” OR “senior” OR “development” OR “developmental” OR “aging” OR “life span”).* The above process yielded 3,484 articles. In addition, 111 studies on young to middle-aged adults from a previous meta-analysis study^21^ were included in the literature pool, 40 of which were excluded according to the current literature exclusion criteria (see below). Moreover, we screened 16 articles citing or being cited by the crucial literature. After removing duplicates, the literature search identified 2,930 articles.

#### Exclusion Criteria

We excluded any articles that met one or more of the following predefined exclusion criteria^89^: 1) not in English or Chinese; 2) not including healthy human participants; 3) case study; 4) not empirical study; 5) not functional resonance imaging (fMRI) or positron emission tomography (PET) study; 6) not whole-brain results (i.e., not have covered the whole gray matter); 7) not in Talairach or Montreal Neurological Institute (MNI) space; 8) not reflecting the congruency effect (i.e., contrasts between incongruent and congruent or between incongruent and neutral conditions); 9) not reporting exact mean age of participants.

A total of 119 articles were identified as eligible for inclusion in our meta-analyses. No statistical methods were used to pre-determine sample sizes, but our sample sizes are similar to or larger than those reported in previous publications^31,90^. Supplementary Fig. S1 shows the preferred reporting items for systematic reviews and meta-analyses (PRISMA)^91^ flow chart for the literature screening process. The 119 articles included 129 studies (individual contrasts reported in the articles) with 3,388 participants and 1,579 activation foci reported. All studies were published or completed between 1994 and 2022. None of the experiments share the same group of participants. The included studies are written in English (124 studies) and Chinese (5 studies). Of the studies included, 125 were published in peer-reviewed journals, and 4 were master’s theses. All included studies reported the task type used, including 74 studies utilizing Stroop-like task (57%), 25 studies utilizing Simon task (19%), 25 studies utilizing Flanker task (19%), 2 studies utilizing a combination of Simon and Flanker tasks (2%), 1 study utilizing a combination of Simon and Stroop tasks (1%), and 3 studies utilizing multisource interference task (2%). In addition, the contrasts conducted to reveal brain activations were also reported, with 98 studies (76%) resulting from the contrast of Incongruent trials > Congruent trials, 25 studies (19%) resulting from the contrast of Incongruent trials > Neutral trials, and 6 studies (5%) resulting from the union contrast of Incongruent trials > Congruent trials and Incongruent trials > Neutral trials. Regarding the handedness of participants in the included studies, 89 studies (69%) included righthanded participants only, 6 studies (5%) included both left and right-handed participants, while 34 studies (26%) did not report this information. Furthermore, 78 studies (60%) included only correct response trials, 2 studies (2%) included both correct and incorrect response trials, while 49 studies (38%) did not report this information. A detailed description of these features for each study is available in the Supplementary Table S1. To eliminate the influence of these confounding factors, we included them as covariates in the modeling analyses.

#### Coding Procedure

A coding manual was formulated to record pertinent study information, including authors, publication dates, experimental tasks, contrasts, and sample demographics (such as the average age and sample size). To ensure coding accuracy, two authors independently coded all studies, with discrepancies resolved through discussion or reference to the original studies. In instances where studies lacked essential information, such as peak coordinates for relevant contrasts, participant age averages, or data for specific age groups, efforts were made to contact the authors via e-mail to obtain the relevant data. In addition, both coordinates and effect sizes (i.e., Hedge’s *g*) were extracted from each study. Further, Talairach space coordinates were transformed to MNI coordinates using the Lancaster transform^92^.

## Meta-Analytic Procedure

### Activation Likelihood Estimation (ALE)

In order to obtain a comprehensive understanding of cognitive control-related brain activity across all age groups and to replicate a prior study^21^, we initially conducted a single dataset meta-analysis using BrainMap GingerALE software^93^ (version 3.0.2, http://www.brainmap.org). This meta-analytical approach, known as activation likelihood estimation, utilizes the spatial convergence of brain activity across multiple studies to determine the probability of activation in specific regions. Foci from individual studies were transformed into a standardized coordinate space and modeled as Gaussian probability values that accounted for variability in the number of participants in each study. In situations where foci overlapped across studies, multiple Gaussians were associated with a single focus, and ALE selected the Gaussian with maximum probability for each focus^93^. Subsequently, ALE score maps were generated by comparing these modeled Gaussian distributions with a null distribution that simulated random brain effects. The null distribution was generated using the same sample size and number of foci groups as the experimental dataset for 1,000 times^94^. ALE scores were then used to calculate *p*-values, which were based on the proportion of values higher than a certain threshold in the null distribution. This resulted in a statistical ALE map that differentiated true brain effects from random effects. A cluster-defining threshold of *p* < 0.001 and a minimum cluster size of 10 voxels (80 mm^3^) were utilized to compute ALE maps, consistent with the threshold applied in the seed-based *d* mapping (SDM) approach (see below).

### Seed-based *d* (Effect Size) Mapping

SDM is an alternative approach to statistically synthesize results from multiple neuroimaging experiments^86^. Similar to ALE, SDM employs a coordinate-based randomeffect approach to amalgamate peak coordinate information into a standard space across several experiments. However, while ALE solely considers the binary feature (i.e., active versus inactive) of peak coordinates, SDM takes into account the quantitative effect size (can be positive or negative) connected to each peak and reconstructs the initial parametric maps of individual experiments before amalgamating them into a meta-analytic map^95^. Therefore, the use of a distinct algorithm in SDM from ALE allows us to scrutinize the robustness and replicability of the outcomes obtained via ALE. More importantly, SDM enables the inclusion of covariates in the meta-regression analyses to reflect the changes in brain function across the lifespan.

We conducted three types of analyses using the SDM approach. Firstly, we estimated the mean activation across all age groups and compared the results with ALE’s single dataset meta-analysis results. Secondly, we conducted contrasts between two groups of studies (e.g., between young to middle-aged adults and a combination of the youth and elderly groups) to identify brain regions that showed different levels of activity across age. This analysis method served to investigate the hypothesized lifespan trajectories, such as the inverted U-shaped pattern by elucidating neural activity variations linked to age. Thirdly, we defined specific models (e.g., linear and square root models) to fit the whole brain to validate brain regions adhering to the hypothetical lifespan changing patterns. This type of analysis aimed to explore various lifespan trajectories, recognizing that different brain regions might follow distinct model functions. See below for the details.

These analyses were conducted using the software of SDM with permutation of subject images (SDM-PSI) (version 6.22, https://www.sdmproject.com). Effect size maps were built for the 129 individual experiments. This was accomplished by (a) converting the statistical value of each peak coordinate into an estimate of effect size (Hedge’s *g*) using standard formulas^96^ and (b) convolving these peaks with a fully anisotropic unnormalized Gaussian kernel (*α* = 1, FWHM = 20 mm) within the boundaries of a gray matter template (voxel size = 2×2×2 mm^3^). Imputation (50 times) was conducted for each study separately to obtain a reliable estimate of brain activation maps^95^. In addition, the individual effect size maps were combined using a random-effect general linear model. To assess the statistical significance of activations in the resulting meta-analytic effect size map, 1,000 random permutations of activation peaks within the gray matter template were compared. Finally, the meta-analytic maps were thresholded using a voxel-wise family-wise error (FWE) corrected threshold of *p* < 0.001 and a cluster-wise extent threshold of 10 voxels^97^.

#### Mean Analyses Across all Studies

This analysis aimed to characterize the activation distributions of cognitive control-related brain regions across all studies, which was conducted utilizing the “Mean” function in SDM-PSI software. In order to verify the reliability of the SDM analysis results, we compared the results with the single dataset meta-analysis using ALE. Results from this analysis were further used as regions of interest (ROIs) in the subsequent model fitting analyses (see section “*Generalized Additive Model (GAM) Fitting*” below). The possibility of publication bias for resultant clusters was examined using Egger’s test^98^, in which any result showing *p* < 0.05 was regarded as having significant publication bias. Heterogeneity was evaluated using the *I*^2^ index, which quantifies the proportion of total variability attributable to heterogeneity between studies. A value less than 25% indicates low heterogeneity among the included studies^99^.

#### Contrast Analyses

To test our hypothesis that cognitive control related brain activities follow an inverted U-shaped trajectory with age, we categorized each study based on the mean age of participants into youth (< 18 years), young to middle-aged adults (18−59 years), and elderly (>= 60 years) groups. The age boundaries were determined to minimize age distribution overlap. We utilized SDM-PSI to perform a contrast analysis between the group of young to middle-aged adults and the combination of other groups in order to examine whether there are brain regions that exhibit an inverted U-shaped lifespan trajectory. This was achieved by assigning studies from the young to middle-aged adult group as 1 and all other studies as −1. This analysis yielded two results, one showing higher activity in young to middle-aged adults than the youth and elderly groups, and the other showing the opposite. Like the mean analysis, results from this analysis were used as ROIs in the subsequent model fitting analyses (see sections “Generalized Additive Model (GAM) Fitting” and “Model Simplification and Model Comparison”).

In addition, to explore other possible trajectories, such as the increase-and-stable pattern^5^, we conducted contrast analyses between the youth and the combined group of young to middle-aged and the elderly, as well as between the elderly and the combined group of youth and young to middle-aged adults. Furthermore, to address the controversies in previous studies, we conducted contrast analyses between older and young to middle-aged adult groups, and between the youth and young to middle-aged adult groups, respectively.

#### Meta-regression Analyses

To better describe the possible lifespan trajectories of the whole brain, we carried out meta-regression analyses with the age and/or its derivatives as regressors. Two regressions were conducted across the whole-brain, including a linear regression (with only age as the regressor) and a square root regression (with age and its square root as separate regressors). The linear regression aimed to test regions with increasing/decreasing activity with age, and the square root regression aimed to test regions with the inverted U-shaped trajectories based on the model fitting analyses (see below).

## Data Extraction

Masks were generated for each ROI derived from the mean and contrast analyses in SDM-PSI as described above. Subsequently, we extracted the effect sizes for each mask. Fifty values were obtained from the SDM iterations and subsequently averaged for each study in each region. The iterated variances were also averaged in a similar way. Additionally, we removed the outliers (beyond 3 standard deviations from the mean) in the following model fitting analyses.

## Generalized Additive Model (GAM) Fitting

To precisely estimate the inverted U-shaped trajectories, we adopted the GAM to fit the curves. The GAM allows for flexible, nonparametric smoothing of predictor variables^100^, and has been widely used to depict the lifespan trajectories^101,102^. We implemented GAMs using the “mgcv” package^100^ in R. For each ROI, we fitted a GAM with the following formula:

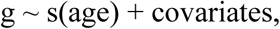

where g is the effect size (dependent variable), s(age) represents a smoothing spline of age (predictor variable), and covariates represent the dummy-coded categorical covariate regressors. These regressors correspond to six aspects of the included studies: (1) the presence of various conflict types (e.g., Stroop or Simon), (2) the mixed subject samples based on handedness (e.g., right handed only or both handed), (3) the different contrasts in reporting congruency effects (e.g., incongruent – congruent or incongruent – neutral), (4) different trial types regarding whether they excluded error trials, (5) the use of different types of experimental design (i.e., event-related or block designs), and (6) the behavioral congruency effects measured by reaction time. Notably, we adopted median imputation^103^ for 9 studies (accounting for 6.98% of the total included studies) not reporting the behavioral congruency effects, and included an indicator regressor to account for the potential impact of imputation^104^. The validity of this imputation approach was confirmed through a robustness analysis (Supplementary Note S4). We incorporated these covariates to control for their potential confounding effects related to age, which could otherwise influence our results. We also adjusted the estimate with a weight parameter, which was the reciprocal of variance. We used penalized regression splines, with the amount of smoothing determined automatically based on generalized cross validation.

For each ROI, we quantified the peak age by choosing the highest prediction of a fine-grained age scale (1,000 points from 8 to 74 years old). We also calculated the estimated degree of freedom (EDF) for the smooth curve by summing up the degree of freedom for each penalized term (i.e., s(age).1 to s(age).9).

## Model Simplification and Model Comparison

While the GAM analysis may yield good fitting results on the data, it is important to acknowledge its potential limitations. One concern is that it can fit the data with high degree of freedoms (up to 7.0, Supplementary Table S4), which makes it susceptible to over-fitting and harder to generalize. Another issue is its poor interpretability. Therefore, we next sought to fit the data with simpler models.

To this end, we used the “metafor” package in R to fit these effect sizes with the age and its derivatives as predictors. Specifically, we tested four non-linear models (see below formulas and Fig. 3). Quadratic and cubic models were included based on previous studies^46,47^; quadratic logarithmic and square-root models were included to capture the possible skewed trajectory, which would reflect the asymmetric trajectory of development and decline of cognitive control^5^. Each model was fitted to each ROI separately. In addition, we included the same covariates as the GAM analysis in each regression model. We calculated the Akaike information criterion (AIC) to evaluate the goodness of fit for each model.

1. Quadratic model *g* ∼ age + age^2^ + covariates
2. Cubic model *g* ∼ age + age^2^ + age^3^ + covariates
3. Quadratic logarithmic model *g* ∼ log(age) + (log(age))^2^ + covariates
4. Square root model *g* ∼ age + sqrt(age) + covariates

## Neurosynth Decoding Analysis

To investigate the potential functional difference between the two sets of brain regions reported in section “*Dissociated Brain Networks with Distinct Lifespan Trajectories*” of Results, we generated two binary maps from the contrast analysis between young to middle-aged adults and the combination of other groups, and then submitted them to the Neurosynth decoding system^41^. As the non-U-shaped map is defined as the converse of the inverted U-shaped map, they were obtained by applying thresholds of 0.1 < *p* < 0.9 and *p* < 0.05, respectively. These lenient threshold boundaries were used to minimize the influence of sparsity on the decoding results. The two-boundary threshold was used in generating the non-U-shaped map, as the original statistical map from the SDMPSI analysis was one-tailed, which means the threshold of *p* > 0.9 indicates the upright U-shaped trend instead of a non-U-shaped trajectory. Additionally, to focus on the brain regions specifically related to cognitive control, the two maps were masked by using results from the mean SDM analysis (Fig. 1B). In the decoding results, we deleted terms that were related to brain regions (e.g., “frontal”), not functional specific (e.g., “task”), and duplicated (e.g., “attention” was removed if there was already “attentional”).

## Laterality Analysis

We calculated the laterality based on the effect size of reported brain coordinates from each study. We computed the sum of effect sizes across coordinates for the left and right hemisphere, respectively, yielding one global effect size each (i.e., *g*_L_ and *g*_R_). Then, we calculated the index of brain laterality with the following equation^105^:

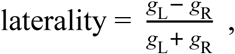

which was then submitted to the GAM and simplified models (i.e., the quadratic, cubic, square root, and quadratic logarithmic models).

To illustrate the contribution of different brain regions to the age-related change of laterality, we calculated the laterality for each voxel^78^. We first used the SDM-PSI to conduct a mean analysis for each age group (i.e., the youth, young to middle-aged adult and elderly), yielding three z-maps. Then the laterality was computed with the above equation for each voxel from the left hemisphere. The opposite values were calculated for the right hemisphere. To avoid the bias due to asymmetric brain hemispheres, we removed voxels without corresponding mirror coordinates. The results were visualized confining to the brain regions estimated from the grand mean analyses (Fig. 1B).

## Data Availability

The meta-data that support the findings of this study are available in Zenodo with the identifier doi: 10.5281/zenodo.12727621106.

## Code Availability

All codes conducting the ROI analyses are available in Zenodo with the identifier doi: 10.5281/zenodo.12727621106.

## Acknowledgement

We thank Fergus I.M. Craik, Beatriz Luna, and Haiyan Wu for valuable suggestions on this manuscript. We also thank Jing Yang and Yifan Zhang for the data check of extracted coordinates. Z.L. discloses support for the research of this work from Scientific Research Fund of Zhejiang Provincial Education Department (Y202249966), Starting Research Fund from Hangzhou Normal University (2021QDL079), and STI 2030-Major Projects (2021ZD0201705). I.T.P discloses support for the research of this work from the Eunice Kennedy Shriver National Institute of Child Health and Human Development (NICHD) (HD098235). G.Y. discloses support for the research of this work from Jiefeng Jiang, and China Postdoctoral Science Foundation (2019M650884). X.L. discloses support for the research of this work from the Ministry of Science and Technology of the People’s Republic of China (2021ZD0200505). The funders had no role in study design, data collection and analysis, decision to publish or preparation of the manuscript.

## Author Contributions Statement

Conceptualization: Z.L. and G.Y.; Methodology: G.Y.; Formal analysis: G.Y. and Z.L. ; Data curation: Z.L.; Writing original draft: G.Y. and Z.L.; Writing, review, and editing: G.Y., Z.L., I.T.P., L.W., X.L. and J.R.; Funding: Z.L., I.T.P., X.L. and G.Y; Supervision: G.Y.

## Inclusion & Ethics

All collaborators of this study fulfilled the criteria for authorship required by Nature Portfolio journals have been included as authors, as their participation was essential for implementation of the study and improvement of the manuscript. Roles and responsibilities were agreed among collaborators ahead of the research. This work does not include findings that are locally relevant. As a meta-analysis, this study does not involve direct interactions with human participants or the collection of new data. Each of the original studies included in this meta-analysis obtained ethical approval from their respective institutions, and we confirm that we complied with all applicable ethical guidelines and regulations.

