## Supplementary material for "The Lifespan Trajectories of Brain Activities Related to Cognitive Control"

### Supplementary Notes

#### Note S1. Validation of Using Mean Age as a Predictor in Meta-regression

Previous meta-regression analyses have typically utilized averaged variables from each included study as predictors[1-3]. A potential concern with this approach is whether the average variable can effectively represent the entire sample, particularly when the range is extensive. In our study, the age ranges for most (~90%) studies are within 30 years, whereas some studies on young to middle-aged adults exhibited wider ranges (up to 59 years). To examine the efficacy of using mean age as a predictor in the meta-regression, we conducted a simulation analysis.

The logic behind this simulation parallels the seed-based *d* mapping with permutation of subject images (SDM-PSI) method[4], which estimates the overall effect size by simulating individual whole-brain activities based on the coordinate effect size data at the study level. By utilizing the average effect size and variance of both the age predictor and brain activity data, we generated simulated individual data to capture a spectrum of possibilities present in the original datasets. Through substantial repetitions, the average results from these individual simulations aimed to provide a reliable approximation of the actual results.

For each study, we generated a set of (N = sample size) individual data points from a normal distribution, defined by the study's mean and standard deviation (square root of variance). This process was conducted for both age and effect size. Simulated data for both variables were constrained within three standard deviations of the mean, with the age further constrained within reported ranges. When age ranges were not available, we used the following specific age brackets: 0 to 17 years for youth studies, 18 to 59 years for young to middle-aged adult studies, and 60 to 100 years for older adult studies. The latter constraint ensured that the simulated ages adhered to the respective age group definitions. The resulting simulated age data was used to predict the simulated effect size data through a mixed-effect generalized additive model (GAM), with both the intercept and age predictor as random effects at the study level. To focus on the age effect, we did not include other covariates such as the task types.

The estimates derived from the mixed-effect GAM were then employed to predict the effect size corresponding to each study's average age value. We correlated these predicted trajectories from the simulation with those from the group-level GAM meta-regression reported in the main text. The simulation process was repeated 1,000 times to ensure robustness. The average correlation across all iterations was interpreted as a measure of overall efficacy.

We applied the simulation test on the nine regions of interest (ROIs) identified from the mean analysis, as these regions yielded more variance in their trajectory shapes. This diversity in the test data could improve the generalizability of our validation approach across different scenarios. Results suggested that, for 8 out of the 9 ROIs, the correlations between the trajectories predicted by the meta-regression and those from the simulated individual data were over 0.96, indicating a substantially high level of agreement. The only deviation was observed in the right inferior temporal gyrus, where the correlation was a respectable 0.73. In this region, the meta-regression produced a linear trend (albeit not significant, as shown in Fig. 2), in contrast to the non-linear trend estimated in the simulated individual data. This discrepancy is likely due to the broader age ranges covered by the simulated data, which could lead to larger variance within a narrow age range. Visual comparisons of the trajectories further highlight strong consistencies between the simulation results and the meta-regression analyses (Supplementary Fig. S4).

Overall, the simulation results indicate that meta-regression, which employs average age as a representative measure of the age range, accurately captures the age-related change patterns that are otherwise estimated with individual data.

**Note S2. Testing the Significance of Peak Ages**

With Simonsohn's[5] two-line approach, we tested whether the visually inverted U-shaped trajectories could be translated into two significant linear trends with opposite directions. We divided the data points into two sets (i.e., pre-peak and after-peak sets) with the peak age as the breaking point, and then conducted linear meta-regression analyses on each set. Given our specific hypothesis of an increasing slope before the peak and a decreasing slope afterward, we adopted one-tailed tests. The resulting p values were false discovery rate (FDR) corrected across the two sets and across the identified brain regions. We applied the same test to the peak ages from both the GAM and the simplified model fitting analyses.

Regarding regions identified in the mean analysis, significant linear relationships were observed for both pre- and post-peak sections in three of the four regions exhibiting significant GAM results (Fig. 2, panels r-IFG, l-ITG, and r-CN), with *p*-FDRs $< 0.05$. The other region (Fig. 2, panel l-ACC) showed significant linear effects in the pre-peak phase, with *p*-FDRs $= 0.006$, but not in the post-peak phase, with *p*-FDRs $= 0.190$.

For the regions identified from the contrast analysis (Fig. 4), the peak ages calculated from the GAM analyses yielded significant linear relationships for both the pre-peak and post-peak sets, *p*-FDRs $< 0.05$, except the right IPL, which only showed pre-peak linearly increase, *p*-FDR $< 0.001$, but not post-peak, *p*-FDR $= 0.053$. Similarly, peak ages identified from the simplified modeling analyses yielded significant linear relationships for most of the regions, with *p*-FDRs $< 0.05$ for both pre- and post-peak sets; the only exception was the left inferior frontal gyrus, which showed marginal significance in the post-peak sets, $p$-FDR = 0.078. Note that while we did not explicitly label the peak in this region due to its marginal statistical significance, we considered it as the potential peak candidate and included it in our main text analysis that correlated peaks between GAM and simplified modeling.

For the laterality analyses (Fig. 7), the peak age calculated from the GAM analysis yielded insignificant linear relationships for both the pre-peak and post-peak sets, $p$s > 0.38, uncorrected. Similarly, peak ages identified from the square root model yielded insignificant linear relationships, $p$s > 0.13, uncorrected.

**Note S3. Contrast Results: Young to middle-aged Adults vs. the Youth and Young to middle-aged Adults vs. the Elderly**

With direct contrast analyses, we found that the young to middle-aged adult group showed significantly higher activation in certain regions compared to the youth group, including bilateral inferior frontal gyrus, right supramarginal gyrus, left inferior parietal lobule, right supplementary motor area, and bilateral insula. No significant regions showed a higher activation in the youth group than the young to middle-aged adult group (Supplementary Fig. S6 and Table S9). Similarly, regions significantly more activated in the young to middle-aged adult group compared to the elderly group were revealed, including right inferior frontal gyrus, left insula, bilateral anterior thalamic projections, right supramarginal gyrus, left supplementary motor area, right inferior parietal lobule, and left caudate nucleus. No significant regions showed higher activity in the elderly group than the young to middle-aged adult group (Supplementary Fig. S8 and Table S9). In addition, there were no significant regions observed when comparing the youth group to the elderly group in either direction (Supplementary Table S9).

In addition, we tested whether behavioral performance might influence the relative brain activity among the three groups. We conducted similar contrast analyses, but adding the covariate of the behavioral congruency effect measured by reaction time, along with an indicator regressor[6-8]. This analysis revealed consistent results as outlined above: young to middle-aged adults exhibited greater activation than the youth in the bilateral inferior frontal gyrus, right supramarginal gyrus, right supplementary motor area, left inferior parietal lobule, bilateral insula, and right inferior temporal gyrus, with no regions showing the opposite (Supplementary Fig. S7 and Table S9). Similarly, young to middle-aged adults showed stronger activation than the elderly in right inferior frontal gyrus, left insula, bilateral anterior thalamic projections, right supramarginal gyrus, left supplementary motor area, bilateral inferior parietal lobule, and left caudate nucleus, with no regions showing the opposite (Supplementary Fig. S9 and Table S9). In addition, the youth showed no difference with the elderly. Overall, the consistency of these findings with the results before controlling for behavioral performance suggests that the observed weaker brain activations in the elderly and youth, compared to young to middle-aged adults, are robust.

In addition, we also conducted similar contrast analyses to the laterality measurement. This was achieved with linear meta-regression models: one comparing the youth with young to middle-aged adults, and the other comparing the elderly with young to middle-aged adults. In each model, we introduced a variable that encoded the specific groups of studies involved, designating the youth and elderly groups with a code of 1, and the young to middle-aged adult group with a code of −1. We also included covariates as other analyses to control for potential confounding factors, such as variations in task types. Results revealed that both the youth ($z = 2.25$, $p = 0.012$, one-tailed, $b = 0.22$, 95% CI = [0.06, 0.38]) and the elderly ($z = 2.06$, $p = 0.020$, one-tailed, $b = 0.20$, 95% CI = [0.04, 0.36]) groups exhibited greater left lateralization than the young to middle-aged adult group.

**Note S4. Robustness analyses**

We conducted several robustness analyses.

In our main text, we employed median imputation to handle missing data related to behavioral congruency effects. However, this approach may introduce bias if the imputed median values differ significantly from the true data. To assess its potential impact on our conclusions, we compared results obtained after removing these studies and those obtained using median imputation for the behavioral congruency effects. We focused on testing two key findings. The first was the inverted U-shaped trajectories observed between young to middle-aged adults and the other groups. To test this, we reanalyzed the fitting process for brain regions identified in the contrast analysis of adult versus other groups (Fig. 4) by removing the 9 studies with missing data. The results showed that the GAM significantly fit all regions with smooth curves, $F$s > 4.1, $p$s < 0.01, FDR corrected, with degrees of freedom varying from 3.1 to 6.6. Using simplified models, we found that the square root model continued to best describe the data, including the left inferior parietal lobule, which had previously fit better with the quadratic model. These results closely replicated the results using imputation. The second finding was greater left lateralization in the youth and elderly compared with the young to middle-aged group. We removed studies with missing data and re-analyzed the laterality data. The results confirmed our original findings: laterality followed a U-shaped trajectory, F(2.9, 3.7) = 3.85, $p$ = 0.008, $R^2$ = 0.22; simplified model fitting suggested that a square root function provided the best goodness of fit, with $\beta_{\text{sqrt(age)}}$ = −0.72 (95% CI = [−1.09, −0.35]), $p$ < 0.001; group analysis showed that both the youth

($z = 2.14$, $p = 0.016$, one-tailed, $b = 0.23$, 95% CI = [0.02, 0.43]) and the elderly ($z = 2.19$, $p = 0.014$, one-tailed, $b = 0.23$, 95% CI = [0.02, 0.43]) exhibited greater left lateralization than the young to middle-aged adult group. These results suggest that our findings based on the imputation method are robust.

It has been suggested that the inclusion of unpublished and non-English studies in meta-analyses can reduce potential bias[10,11]. However, this practice may also introduce variations in the results. Our study incorporated four unpublished master's theses and five non-English studies written in Chinese. To test their influence, we performed a meta-regression analysis after excluding these studies, using regions identified by the mean analysis of SDM. Given that all four unpublished studies were also non-English (a total of five), we excluded both from the reanalysis due to significant overlap. Results showed that GAM significantly fitted the age-related changes in activation levels for 4 out of the 9 regions, *p*s < .05, FDR corrected (Supplementary Fig. S5, l-ACC, r-IFG, l-ITG, and r-CN), and the other five regions showed no significant age-related changes. These results closely replicate the original findings. Thus, the inclusion of unpublished and non-English studies appears to have a limited impact on our meta-analysis results.

Furthermore, to assess the potential impact of missing studies within the 45-60 age range on the mean results, we conducted a mean analysis including the age as a covariate using the SDM. This analysis reaffirmed the same brain clusters previously identified (see Supplementary Fig. S3 and Table S3). Further examination revealed an overlap of 21,144 voxels between the two results, which constitutes 85.9% of their combined total of 24,620 voxels. This suggests that the missing studies have limited influence on the mean results.

### Supplementary Discussion

#### Implications of Our Finding for Other Cognitive Control Subdomains

A more comprehensive understanding of the lifespan trajectory of cognitive control relies on studying different subdomains of cognitive control, such as conflict processing, working memory, and cognitive flexibility[12]. Previous studies have suggested that different subdomains of cognitive control exhibit both similar and separable neural mechanisms[13]. Moreover, these subdomains might follow different lifespan trajectories[14]. Findings from our study may contribute valuable insights into the potential differences in brain activity across different components of cognitive control during development.

Consistent with the inverted U-shaped trajectory, we confirmed that the young adults exhibit higher levels of cognitive control brain activities compared to the youth during conflict processing but not otherwise (supplementary Figure S3, Table S9). This observation is in line with previous studies[15-19]. To the best of our knowledge, previous research using conflict tasks has rarely reported findings that contradict this direction except [20]. However, more controversial patterns have emerged in studies focusing on other subdomains of cognitive control, such as working memory[21-23] and response inhibition[24-26]. For instance, with visual-spatial working memory task, Scherf, Sweeney [21] found an age-related increase in the recruitment of brain regions in the lateral prefrontal cortex, whereas Geier, Garver [23] reported both age-related increase and decrease in different prefrontal regions. These studies employed similar paradigms but different contrast methods (i.e., delay vs. non-delay and long-delay vs. short-delay), which exemplifies the many diversities within the working memory domain[27]. Similarly, in the subdomain of response inhibition, studies have shown discrepancies in brain engagement with different designs and different measurements[28,29]. On the contrary, conflict processing provides a more consistent framework for measuring brain activities across age groups. Given these differences, it is prudent to be cautious in generalizing our current findings to the lifespan trajectories of other subdomains of cognitive control, although an inverted U-shaped trajectory could be a good candidate hypothesis to begin with. Future research is necessary to develop more precise cross-age comparisons within these other cognitive control subdomains.

### Supplementary Figures

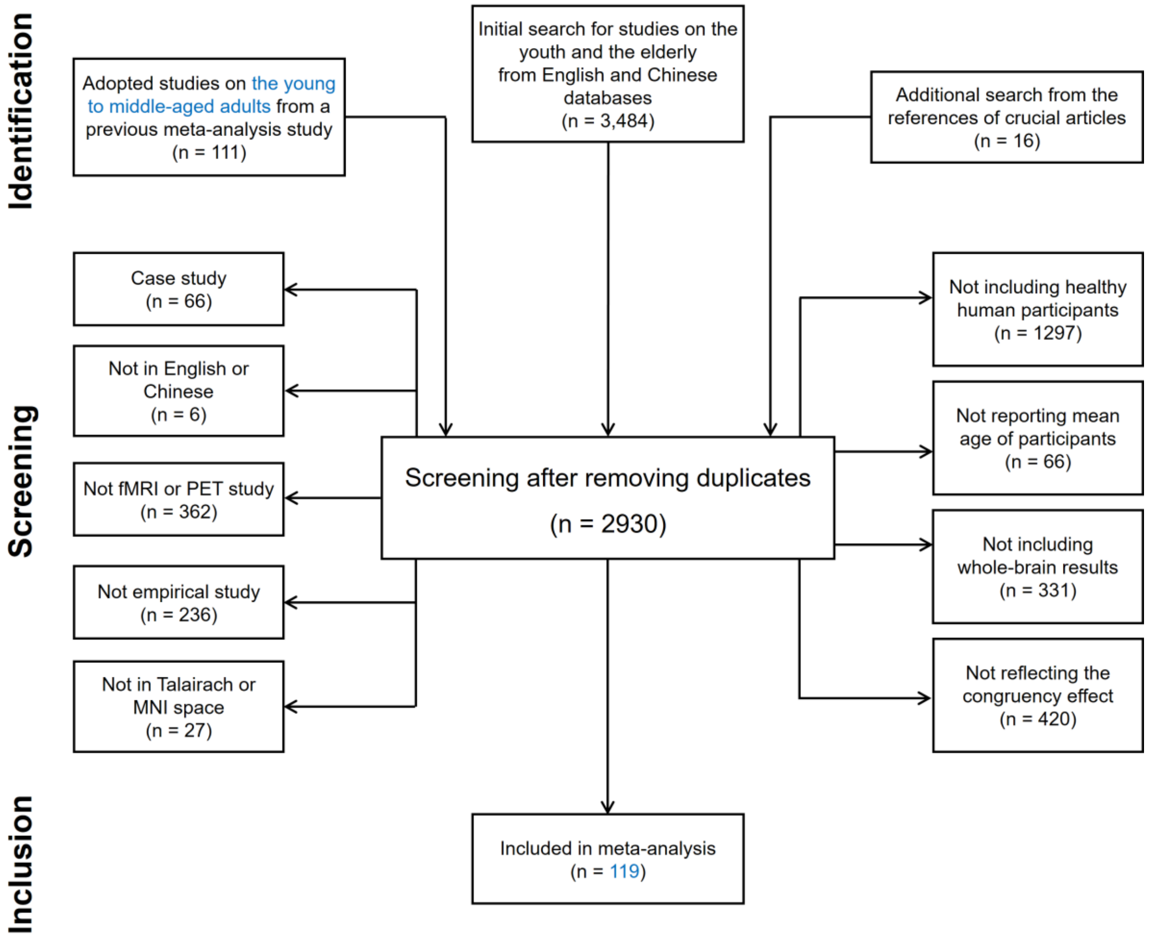


**Fig. S1**. The preferred reporting items for systematic reviews and meta-analyses (PRISMA) flowchart of the selection process for included articles. fMRI = functional magnetic resonance imaging; PET = positron emission tomography; MNI = Montreal Neurological Institute.

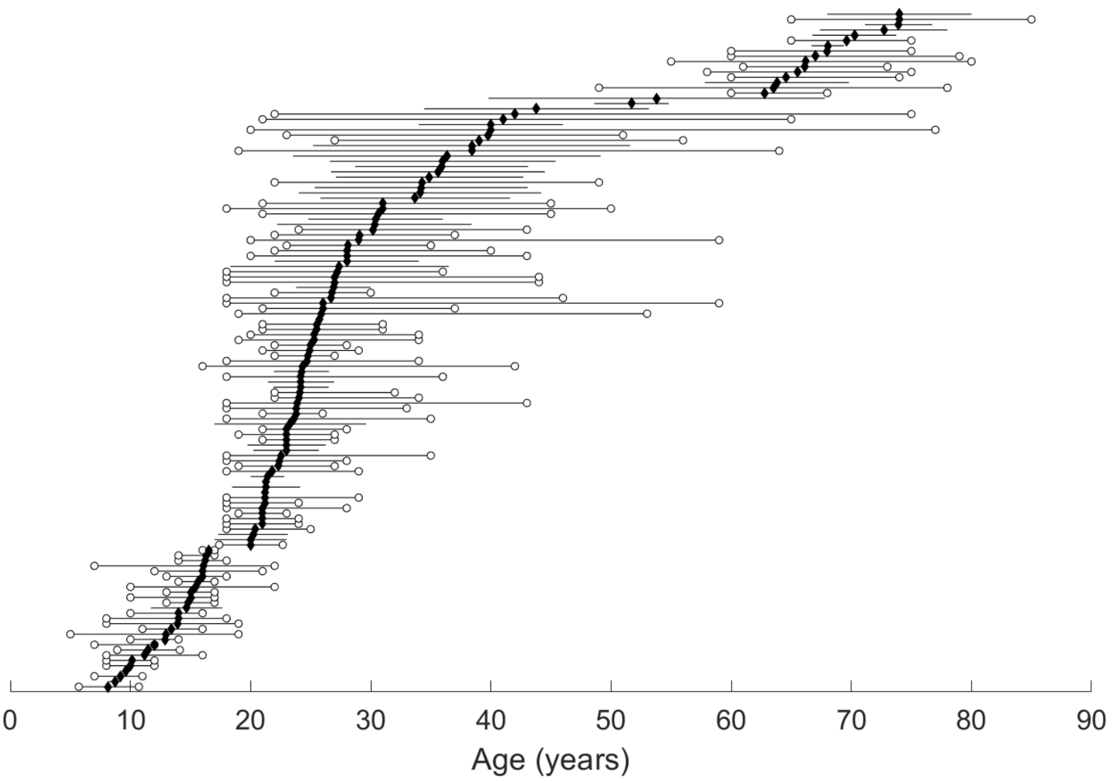


**Fig. S2.** Age distribution of included studies. Black diamonds denote mean ages, lines with open circles denote age ranges, and lines without open circles denote standard deviations. Ages are sorted with the mean age of each study.

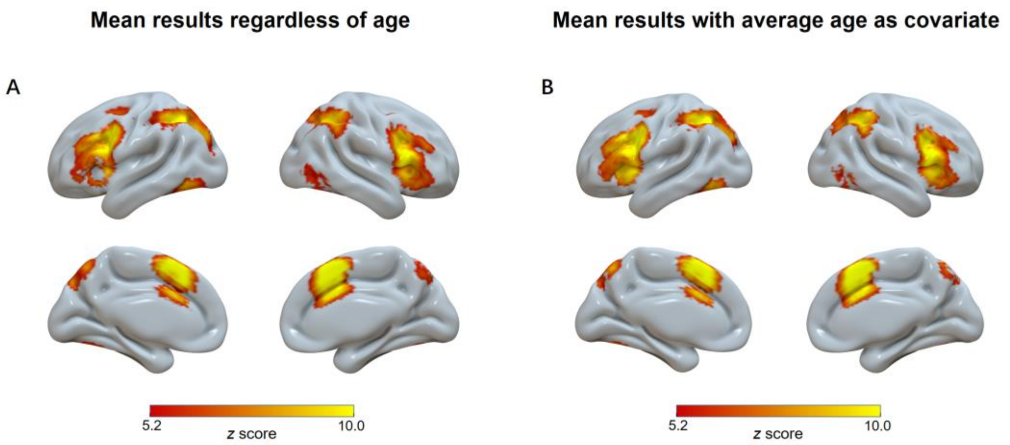


**Fig. S3**. Significant clusters (voxel-wise FWE-corrected, $p < 0.001$, with minimum cluster size ≥ 10 voxels) across all studies in the SDM meta-analysis regardless of age (A) and with average age as covariate (B).

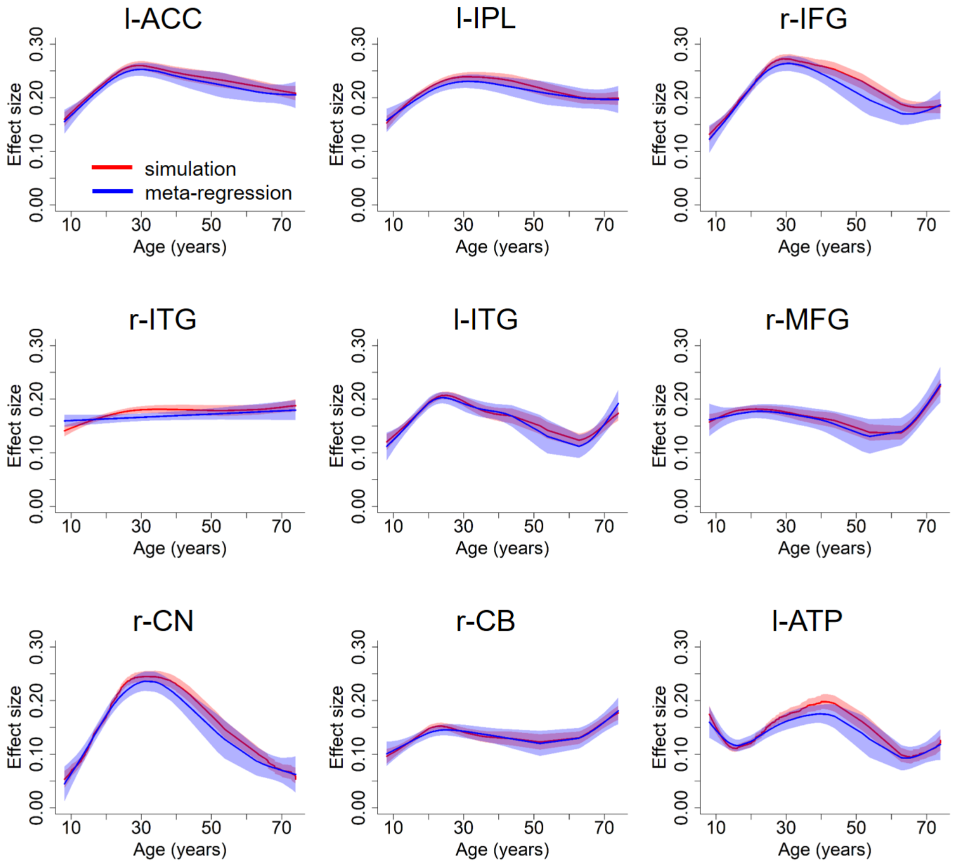


**Fig. S4.** Trajectories predicted by the simulation (red) and meta-regression (blue). The trajectory of meta-regression was consistent with what was reported in Fig. 2 of the main text, except that in panels l-IPL, r-ITG, r-MFG, r-CB and l-ATP, where the fitted curves were not plotted in Fig. 2 due to insignificant GAM fitting results. The simulation predicted curves derive from the simulated individual data, with the illustrated trajectories representing the average curves across 1,000 iterations. Shaded regions around these curves represent standard errors.

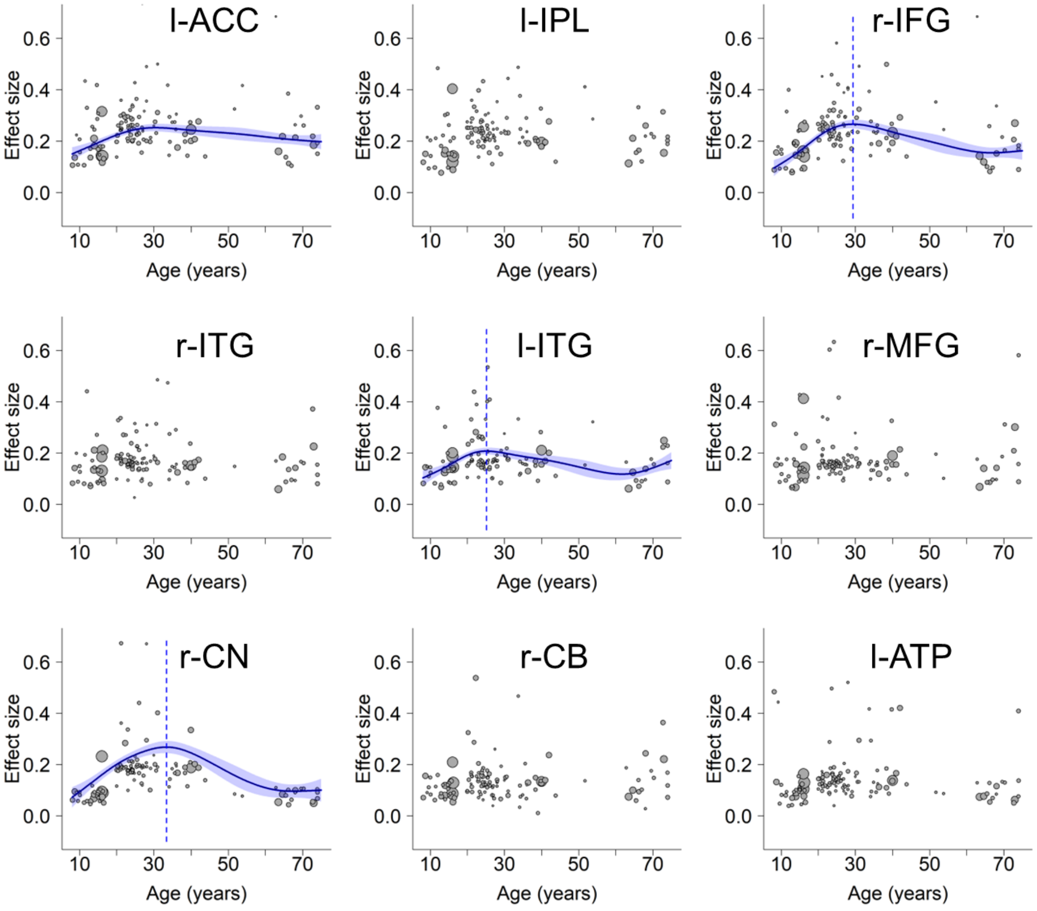


**Fig. S5.** Robustness analysis of lifespan trajectories within regions identified in the mean analysis, with the unpublished and non-English studies excluded. l-ACC: left anterior cingulate cortex, l-IPL: left inferior parietal lobule, r-IFG: right inferior frontal gyrus, r-ITG: right inferior temporal gyrus, l-ITG: left inferior temporal gyrus, r-MFG: right middle frontal gyrus, r-CN: right caudate nucleus, r-CB: right cerebellum, l-ATP: left anterior thalamic projections. Scattered plots are the effect sizes as a function of age, with curves fitted by GAM. The sizes of the scattered dots show the square root of model weights (1/variance) for each study. Shaded areas around the curves represent standard errors. Dashed lines indicate peak ages. L-ACC does not show the peak age due to an insignificant decrease at the later stage.

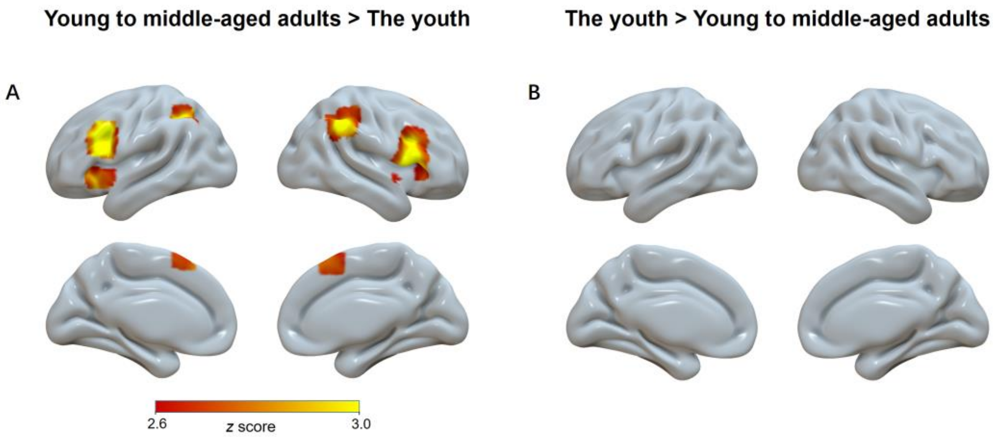


**Fig. S6**. Significant clusters (voxel-wise FWE-corrected, $p < 0.001$, with minimum cluster size ≥ 10 voxels) for contrast analyses between the young to middle-aged adult group and the youth group. The displayed regions show higher activities in young to middle-aged adults than the youth (A). No regions show the opposite direction (B).

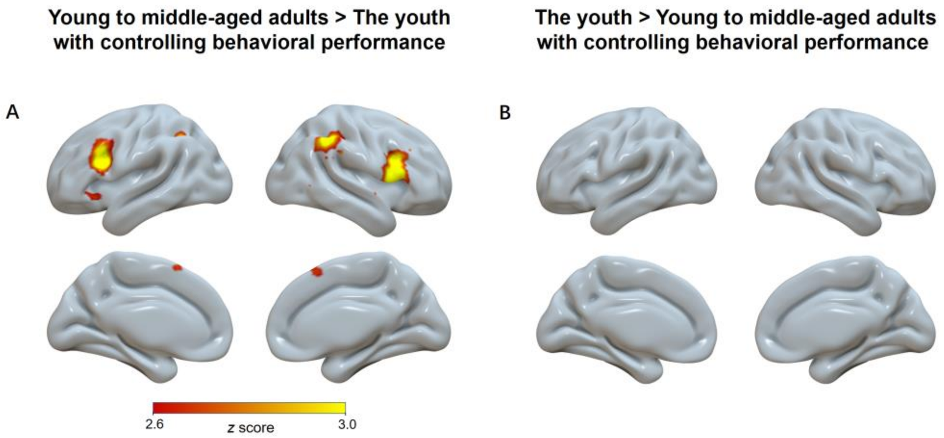


**Fig. S7**. Significant clusters (voxel-wise FWE-corrected, $p < 0.001$, with minimum cluster size ⩾ 10 voxels) for contrast analyses between the young to middle-aged adult group and the youth group with controlling behavioral performance. The displayed regions show higher activities in young to middle-aged adults than the youth (A). No regions show the opposite direction (B).

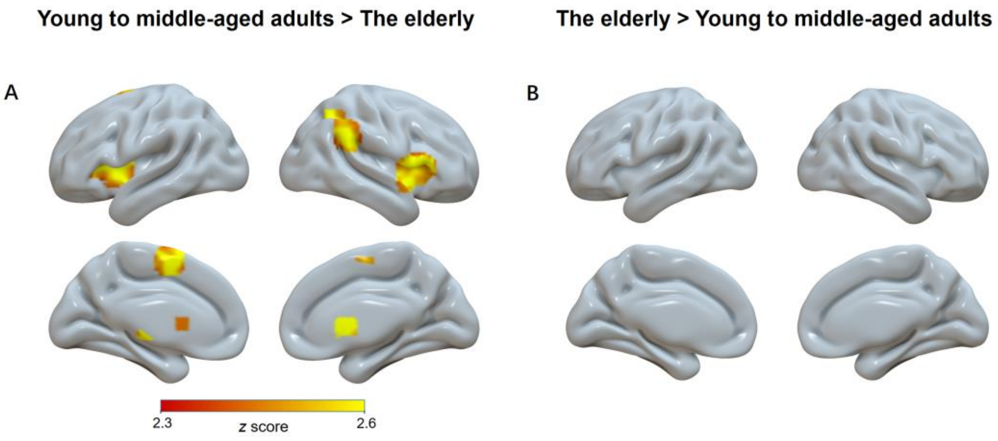


**Fig. S8**. Significant clusters (voxel-wise FWE-corrected, $p < 0.001$, with minimum cluster size ≥ 10 voxels) for contrast analyses between the young to middle-aged adult group and the elderly group. The displayed regions show higher activities in younger adults than older adults (A). No regions show the opposite direction (B).

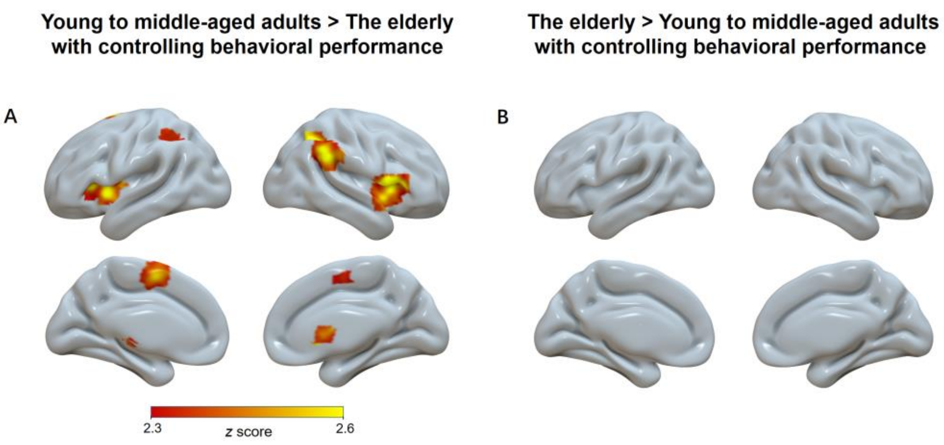


**Fig. S9**. Significant clusters (voxel-wise FWE-corrected, $p < 0.001$, with minimum cluster size ≥ 10 voxels) for contrast analyses between the young to middle-aged adult group and the elderly group with controlling behavioral performance. The displayed regions show higher activities in younger adults than older adults (A). No regions show the opposite direction (B).

### Supplementary Tables

**Table S1**. Studies included in the present study (see the Excel file “Supplementary Table S1”).

**Table S2**. Activation across all age groups with ALE (voxel-wise FWE-corrected, $p <$ 0.001, with minimum cluster size ≥ 10 voxels).

| Order | # Voxels | ALE ($\times 10^{-2}$) | *p* | L/R | MNI coordinate | | | Anatomical location | BA |
|---|---|---|---|---|---|---|---|---|---|
| | | | | | x | y | z | | |
| 1 | 859 | 12.295 | < 0.001 | / | 0 | 14 | 48 | pre-supplementary motor area | 32 |
| 2 | 682 | 6.862 | < 0.001 | L | −30 | −62 | 40 | inferior parietal lobule | 39 |
| 3 | 538 | 9.594 | < 0.001 | L | −42 | 4 | 32 | dorsolateral prefrontal cortex | 6 |
| 4 | 402 | 9.378 | < 0.001 | R | 46 | 10 | 30 | inferior frontal gyrus | 9 |
| 5 | 364 | 9.135 | < 0.001 | R | 36 | 22 | 0 | insula | 13 |
| 6 | 355 | 8.223 | < 0.001 | R | 32 | −52 | 46 | inferior parietal lobule | 7 |
| 7 | 207 | 7.329 | < 0.001 | L | −34 | 20 | 0 | insula | 13 |
| 8 | 143 | 6.356 | < 0.001 | L | −26 | −6 | 52 | frontal eye field | 6 |
| 9 | 58 | 5.448 | < 0.001 | L | −42 | −66 | −10 | inferior temporal gyrus | 37 |
| 10 | 27 | 4.963 | < 0.001 | R | 26 | 0 | 50 | Frontal eye field | 6 |

*Note.* ALE = activation likelihood estimation; MNI = Montreal Neurological Institute; BA = Brodmann area; L = left; R = right.

**Table S3**. Activation across all age groups with SDM (voxel-wise FWE-corrected, $p < 0.001$, with minimum cluster size ≥ 10 voxels).

| Order | # Voxels | *Z* | *p* | L/R | MNI coordinate | | | Anatomical location | BA |
|---|---|---|---|---|---|---|---|---|---|
| | | | | | x | y | z | | |
| **Mean results regardless of age** | | | | | | | | | |
| 1 | 9240 | 12.751 | < 0.001 | L | −2 | 26 | 36 | anterior cingulate cortex | 24 |
| 2 | 6695 | 11.876 | < 0.001 | L | −36 | −62 | 44 | inferior parietal lobule | 7 |
| 3 | 4714 | 12.257 | < 0.001 | R | 50 | 20 | 4 | inferior frontal gyrus | 45 |
| 4 | 1247 | 7.531 | < 0.001 | R | 50 | −62 | −12 | inferior temporal gyrus | 37 |
| 5 | 1075 | 8.977 | < 0.001 | L | −44 | −62 | −12 | inferior temporal gyrus | 37 |
| 6 | 197 | 6.599 | < 0.001 | R | 34 | 4 | 52 | middle frontal gyrus | 6 |
| 7 | 41 | 5.939 | < 0.001 | R | 10 | 4 | 10 | caudate nucleus | / |
| 8 | 20 | 5.827 | < 0.001 | R | 14 | −70 | −22 | cerebellum | 18 |
| 9 | 11 | 5.476 | < 0.001 | L | −12 | −2 | 16 | anterior thalamic projections | / |
| **Mean results with average age as covariate** | | | | | | | | | |
| 1 | 9410 | 12.848 | < 0.001 | L | −2 | 26 | 40 | anterior cingulate cortex | 24 |
| 2 | 6527 | 12.088 | < 0.001 | L | −36 | −62 | 44 | inferior parietal lobule | 7 |
| 3 | 4444 | 12.5 | < 0.001 | R | 50 | 18 | 4 | inferior frontal gyrus | 45 |
| 4 | 1026 | 8.91 | < 0.001 | R | 52 | −62 | −10 | inferior temporal gyrus | 37 |
| 5 | 988 | 9.129 | < 0.001 | L | −46 | −62 | −12 | inferior temporal gyrus | 37 |
| 6 | 75 | 6.275 | < 0.001 | R | 34 | 6 | 50 | middle frontal gyrus | 6 |
| 7 | 19 | 6.034 | < 0.001 | R | 10 | 4 | 10 | caudate nucleus | / |
| 8 | 18 | 6.195 | < 0.001 | L | −12 | 0 | 16 | anterior thalamic projections | / |
| 9 | 17 | 5.754 | < 0.001 | R | 10 | −20 | 2 | anterior thalamic projections | / |

*Note*. MNI = Montreal Neurological Institute; BA = Brodmann area; L = left; R = right.

**Table S4**. The fitting results of the GAM analyses.

| Order | Region name | Estimated degree of freedom | Peak age | F | Effect size |
|---|---|---|---|---|---|
| **Regions identified in the mean analysis with SDM** | | | | | |
| 1 | anterior cingulate cortex | 3.1 | 30.7 | 3.768 | 0.196 |
| 2 | inferior parietal lobule | 2.9 | / | 2.525 | 0.222 |
| 3 | inferior frontal gyrus | 4.0 | 29.6 | 8.202 | 0.252 |
| 4 | inferior temporal gyrus | 1.0 | / | 0.048 | -0.012 |
| 5 | inferior temporal gyrus | 4.0 | 25.3 | 3.637 | 0.114 |
| 6 | middle frontal gyrus | 3.4 | / | 1.514 | 0.326 |
| 7 | caudate nucleus | 3.7 | 33.4 | 8.483 | 0.269 |
| 8 | cerebellum | 3.3 | / | 1.581 | 0.009 |
| 9 | anterior thalamic projections | 5.1 | / | 2.466 | 0.148 |
| **Regions identified in the contrast analysis of Young to middle-aged adults > Others** | | | | | |
| 1 | right inferior frontal gyrus | 4.6 | 30.2 | 14.821 | 0.409 |
| 2 | left inferior frontal gyrus | 4.1 | 30.7 | 8.885 | 0.301 |
| 3 | right inferior parietal lobule | 3.6 | 32.2 | 8.398 | 0.298 |
| 4 | left supplementary motor area | 2.9 | 32.2 | 5.152 | 0.162 |
| 5 | left inferior parietal lobule | 3.4 | 39.4 | 11.627 | 0.338 |
| 6 | right caudate nucleus | 3.7 | 33.5 | 10.513 | 0.332 |
| 7 | left insula | 3.4 | 31.1 | 6.471 | 0.232 |
| 8 | left insula | 3.9 | 29.8 | 6.747 | 0.242 |
| 9 | right middle cingulate cortex | 6.7 | 24.5 | 24.128 | 0.648 |

*Note.* Regions are from the contrast analysis between young to middle-aged adults and other age groups (Table 1). Effect size is measured by the adjusted $R^2$.

**Table S5**. Weights of each model based on AIC values.

| **Order** | **Region name** | **Qua** | **Cub** | **QuaLog** | **Sqrt** |
|---|---|---|---|---|---|
| 1 | right inferior frontal gyrus | 0.216 | 0.103 | 0.254 | 0.427$^{*}$ |
| 2 | left inferior frontal gyrus | 0.372 | 0.067 | 0.149 | 0.411$^{*}$ |
| 3 | right inferior parietal lobule | 0.255 | 0.063 | 0.309 | 0.373$^{*}$ |
| 4 | left supplementary motor area | 0.293 | 0.052 | 0.279 | 0.376$^{*}$ |
| 5 | left inferior parietal lobule | 0.508$^{*}$ | 0.067 | 0.080 | 0.345 |
| 6 | right caudate nucleus | 0.284 | 0.069 | 0.246 | 0.401$^{*}$ |
| 7 | left insula | 0.247 | 0.113 | 0.184 | 0.456$^{*}$ |
| 8 | left insula | 0.229 | 0.081 | 0.321 | 0.369$^{*}$ |
| 9 | right middle cingulate cortex | 0.287 | 0.053 | 0.299 | 0.361$^{*}$ |

*Note.* Regions are from the contrast analysis between young to middle-aged adults and other age groups (Table 1). Qua = quadratic model; Cub = cubic model; QuaLog = quadratic logarithmic model; Sqrt = square root model. * denotes the model with the highest weight.

**Table S6**. The results of the optimal model fitting.

| **Order** | **Region name** | **BA** | **Best model** | **QM** | ***p*** | **Peak age** | **Fixed effect** | **95% Confidence interval** |
|---|---|---|---|---|---|---|---|---|
| 1 | right inferior frontal gyrus | 24 | Square root | 15.879 | < 0.001 | 34.1 | 0.337 | [0.145, 0.530] |
| 2 | left inferior frontal gyrus | 7 | Square root | 19.162 | < 0.001 | 36.0 | 0.335 | [0.142, 0.527] |
| 3 | right inferior parietal lobule | 45 | Square root | 12.153 | 0.007 | 33.7 | 0.240 | [0.049, 0.431] |
| 4 | left supplementary motor area | 37 | Square root | 13.026 | 0.008 | 33.3 | 0.237 | [0.046, 0.429] |
| 5 | left inferior parietal lobule | 37 | Quadratic | 22.443 | < 0.001 | 40.0 | −0.0002 | [−0.0004, −0.0001] |
| 6 | right caudate nucleus | 6 | Square root | 14.548 | 0.002 | 33.6 | 0.282 | [0.092, 0.472] |
| 7 | left insula | / | Square root | 29.461 | < 0.001 | 34.2 | 0.378 | [0.184, 0.572] |
| 8 | left insula | 18 | Square root | 11.185 | 0.007 | 34.2 | 0.239 | [0.047, 0.430] |
| 9 | right middle cingulate cortex | / | Square root | 6.200 | 0.021 | 33.6 | 0.198 | [0.007, 0.388] |

*Note.* Regions are from the contrast analysis between young to middle-aged adults and other age groups (Table 1). QM is the test statistic of the Wald-type test of the model coefficients; *p, fixed effect and 95% confidence interval* are the statistical significance of the sqrt(age) and the $(age)^2$ terms for the Square root and Quadratic models, respectively. BA = Brodmann area.

**Table S7**. Brain areas exhibiting a square root lifespan trajectory (voxel-wise FWE-corrected, $p < 0.001$, with minimum cluster size ≥ 10 voxels).

| **Order** | **# Voxels** | ***Z*** | ***p*** | **L/R** | **MNI coordinate** | | | **Anatomical location** | **BA** |
|---|---|---|---|---|---|---|---|---|---|
| | | | | | **x** | **y** | **z** | | |
| 1 | 653 | 4.670 | < 0.001 | R | 50 | 20 | 2 | inferior frontal gyrus | 45 |
| 2 | 637 | 5.440 | < 0.001 | L | −44 | 16 | 24 | inferior frontal gyrus | 48 |
| 3 | 222 | 3.986 | < 0.001 | R | 40 | −50 | 38 | angular gyrus | / |
| 4 | 97 | 3.749 | < 0.001 | L | −36 | −50 | 42 | inferior parietal lobule | 40 |
| 5 | 36 | 3.123 | < 0.01 | R | 34 | 24 | 6 | insula | 48 |
| 6 | 17 | 3.245 | < 0.001 | R | 10 | 4 | 6 | caudate nucleus | / |
| 7 | 15 | 3.392 | < 0.001 | L | −10 | −18 | 2 | anterior thalamic projections | / |

*Note*. MNI = Montreal Neurological Institute; BA = Brodmann area; L = left; R = right.

**Table S8**. The top-3 decoding terms with the Neurosynth database for the inverted U-shaped and non-inverted U-shaped regions. The three terms of interest (i.e., “attention(al)”, “(cognitive) control” and “monitoring”) are also included.

| Rank | Inverted U-shaped | | Non-inverted U-shaped | |
|---|---|---|---|---|
| | **Term** | **Correlation** | **Term** | **Correlation** |
| 1 | Demands | 0.308 | **Attentional** | 0.240 |
| 2 | Working memory | 0.299 | Working memory | 0.226 |
| 3 | **Cognitive control** | 0.187 | Spatial | 0.207 |
| > 50 | Attentional | 0.100 | Monitoring | 0.090 |
| > 50 | Monitoring | 0.072 | Control | 0.062 |

Note. “Cognitive control” corresponds to the frontoparietal control network, “Attentional” corresponds to the dorsal attention network, and “Monitoring” corresponds to the cingulo-opercular network[30].

**Table S9**. Overview of significant clusters in two-group contrast analyses between the youth, young to middle-aged adult, and the elderly groups (voxel-wise FWE-corrected, $p < 0.001$, with minimum cluster size ≥ 10 voxels) with SDM.

| Order | # Voxels | *Z* | *p* | L/R | MNI coordinate | | | Anatomical location | BA |
|---|---|---|---|---|---|---|---|---|---|
| | | | | | x | y | z | | |
| **Young to middle-aged adults > The youth** | | | | | | | | | |
| 1 | 802 | 4.103 | < 0.001 | R | 54 | 14 | 10 | inferior frontal gyrus | 48 |
| 2 | 523 | 4.146 | < 0.001 | R | 54 | −46 | 42 | supramarginal gyrus | 40 |
| 3 | 495 | 4.609 | < 0.001 | L | −44 | 16 | 24 | inferior frontal gyrus | 48 |
| 4 | 206 | 3.710 | < 0.001 | L | −36 | −54 | 42 | inferior parietal lobule | 40 |
| 5 | 105 | 3.086 | < 0.001 | R | 10 | 16 | 54 | supplementary motor area | / |
| 6 | 73 | 3.150 | < 0.001 | L | −36 | 20 | −8 | insula | 47 |
| 7 | 14 | 2.810 | < 0.001 | R | 32 | 26 | −2 | insula | 47 |
| **The youth > Young to middle-aged adults** | | | | | | | | | |
| None | | | | | | | | | |
| **Young to middle-aged adults > The youth with controlling behavioral performance** | | | | | | | | | |
| 1 | 1151 | 4.260 | < 0.001 | R | 56 | 14 | 12 | inferior frontal gyrus | 44 |
| 2 | 801 | 3.951 | < 0.001 | R | 52 | −44 | 40 | supramarginal gyrus | 40 |
| 3 | 654 | 4.441 | < 0.001 | L | −44 | 16 | 24 | inferior frontal gyrus | 48 |
| 4 | 346 | 3.090 | < 0.001 | R | 10 | 20 | 56 | supplementary motor area | / |
| 5 | 286 | 3.481 | < 0.001 | L | −32 | −56 | 42 | inferior parietal lobule | 7 |
| 6 | 140 | 3.219 | < 0.001 | L | −38 | 24 | −6 | insula | 47 |
| 7 | 129 | 2.968 | < 0.001 | R | 32 | 28 | 0 | insula | 47 |
| 8 | 14 | 2.770 | < 0.001 | R | 52 | −62 | 2 | inferior temporal gyrus | 37 |
| **The youth > Young to middle-aged adults with controlling behavioral performance** | | | | | | | | | |
| None | | | | | | | | | |
| **Young to middle-aged adults > The elderly** | | | | | | | | | |
| 1 | 823 | 4.336 | < 0.001 | R | 50 | 12 | 2 | inferior frontal gyrus | 48 |
| 2 | 373 | 3.269 | < 0.001 | L | −30 | 6 | 0 | insula | 48 |

| | | | | | | | | | |
|---|---|---|---|---|---|---|---|---|---|
| 3 | 339 | 3.480 | < 0.001 | R | 10 | 6 | 4 | anterior thalamic projections | / |
| 4 | 192 | 3.122 | < 0.001 | R | 54 | −46 | 34 | supramarginal gyrus | 40 |
| 5 | 109 | 3.187 | < 0.001 | L | −10 | 2 | 60 | supplementary motor area | 6 |
| 6 | 93 | 3.422 | < 0.001 | R | 30 | −56 | 40 | inferior parietal lobule | 7 |
| 7 | 25 | 2.751 | < 0.001 | L | −8 | −20 | 0 | anterior thalamic projections | / |
| 8 | 23 | 2.770 | < 0.001 | L | −10 | 8 | 4 | caudate nucleus | / |
| **The elderly > Young to middle-aged adults** | | | | | | | | | |
| None | | | | | | | | | |
| **Young to middle-aged adults > The elderly with controlling behavioral performance** | | | | | | | | | |
| 1 | 1009 | 4.502 | < 0.001 | R | 48 | 12 | 0 | inferior frontal gyrus | 48 |
| 2 | 465 | 3.451 | < 0.001 | L | −38 | 24 | 6 | insula | 47 |
| 3 | 374 | 3.256 | < 0.001 | R | 10 | 4 | 4 | anterior thalamic projections | / |
| 4 | 249 | 3.162 | < 0.001 | R | 58 | −44 | 32 | supramarginal gyrus | 48 |
| 5 | 176 | 2.991 | < 0.001 | L | −8 | 2 | 62 | supplementary motor area | 6 |
| 6 | 129 | 3.424 | < 0.001 | R | 32 | −54 | 40 | inferior parietal lobule | 7 |
| 7 | 38 | 2.906 | < 0.001 | L | −10 | −20 | 0 | anterior thalamic projections | / |
| 8 | 34 | 2.602 | < 0.001 | L | −36 | −50 | 44 | inferior parietal lobule | 40 |
| 9 | 27 | 2.708 | < 0.001 | L | −8 | 10 | 4 | caudate nucleus | / |
| 10 | 12 | 2.516 | < 0.001 | L | −42 | −48 | 52 | inferior parietal lobule | 40 |
| **The elderly > Young to middle-aged adults with controlling behavioral performance** | | | | | | | | | |
| None | | | | | | | | | |
| **The youth > The elderly** | | | | | | | | | |
| None | | | | | | | | | |
| **The elderly > The youth** | | | | | | | | | |
| None | | | | | | | | | |
| **The youth > The elderly with controlling behavioral performance** | | | | | | | | | |
| None | | | | | | | | | |
| **The elderly > The youth with controlling behavioral performance** | | | | | | | | | |
| None | | | | | | | | | |

*Note*. MNI = Montreal Neurological Institute; BA = Brodmann area; L = left; R = right.
