## supplementary table S1 for "The Lifespan Trajectories of Brain Activities Related to Cognitive Control"

### **Table S1**. Studies included in the present study.

| Order | First author | Year | Publication type | Task type | Experimental design | Sample size (male) | Language | Mean age (SD) (years) | Age range (years) | Handness | Contrast | Correct trials only | Congruency effect (reaction time, ms) |
|---|---|---|---|---|---|---|---|---|---|---|---|---|---|
| **The youth** | | | | | | | | | | | | | |
| 1 | Andrews-Hanna | 2011 | Journal | Stroop task | Hybrid block/event-related design | 32 (17) | English | 15.6 (*n.r.*) | 14−17 | right | I > N | Yes | 78 |
| 2 | Bernal | 2009 | Journal | Stroop task | Hybrid block/event-related design | 18 (8) | English | 14.67 (2.97) | *n.r.* | right | I > C | *n.r.* | *n.r.* |
| 3 | Bunge | 2002 | Journal | Flanker task | Event-related design | 16 (10) | English | 10 (*n.r.*) | 8−12 | right | I > N | *n.r.* | 44 |
| 4 | Cao | 2006 | Journal | rhyming task (Stroop-like) | Event-related design | 14 (8) | English | 11.5 (*n.r.*) | 8.9−14.11 | right | I > N | No | 121 |
| 5 | Carp | 2012 | Journal | Simon task | Event-related design | 18 (10) | English | 14 (*n.r.*) | 8−18 | *n.r.* | I > C | Yes | 283.3 |
| 6 | de Kieviet | 2014 | Journal | Flanker task | Block design | 47 (21) | English | 8.7 (0.5) | *n.r.* | *n.r.* | I > C | Yes | 150 |
| 7 | Fan | 2014 | Journal | Stroop task | Event-related design | 23 (21) | English | 11.2 (2.9) | 8−16 | *n.r.* | I > C | *n.r.* | 73 |
| 8 | Gee | 2022 | Journal | Stroop task | Block design | 40 (18) | English | 12.93 (3.92) | 5−19 | right | I > C | *n.r.* | 195.04 |
| 9 | Halari | 2009 | Journal | Simon task | Event-related design | 21 (10) | English | 16.3 (1.1) | 14-17 | *n.r.* | I > C | *n.r.* | 94 |
| 10 | Hansen | 2018 | Journal | Stroop task | Block design | 171 (121) | English | 16.2 (1) | 14−18 | *n.r.* | I > C | *n.r.* | *n.r.* |
| 11 | Kaufmann | 2006 | Journal | number-size congruity task (Stroop-like) | Event-related design | 17 (10) | English | 9.6 (*n.r.*) | *n.r.* | right | I > N | Yes | 79 |
| 12 | Kim-Spoon | 2021 | Journal | multi-source interference task | Block design | 151 (*n.r.*) | English | 16 (0.54) | 13−18 | *n.r.* | I > N | Yes | 398.4 |
| 13 | Konrad | 2005 | Journal | Flanker task | Event-related design | 16 (16) | English | 10.1 (*n.r.*) | 8−12 | right | I > C | Yes | 80 |
| 14 | Liu | 2016 | Journal | multi-source interference task | Event-related design | 72 (*n.r.*) | English | 13.9 (3.3) | 8−19 | *n.r.* | I > C | Yes | 314 |
| 15 | Margolis | 2017 | Journal | Simon task | Event-related design | 55 (*n.r.*) | English | 16.1 (3.8) | 7−22 | *n.r.* | I > C | Yes | 40 |
| 16 | Mincic | 2010 | Journal | Stroop task | Event-related design | 35 (18) | English | *n.r.* | 16−17 | right | I > N | *n.r.* | 57.5 |
| 17 | Posner | 2011 | Journal | Stroop task | Block design | 15 (13) | English | 13.4 (1.2) | 11−16 | right | I > N | *n.r.* | 86.6 |
| 18 | Puetz | 2016 | Journal | Stroop task | Block design | 19 (*n.r.*) | English | 12.9 (1.32) | 10−14 | *n.r.* | I > N | *n.r.* | 100 |
| 19 | Rubia | 2006 | Journal | Simon task | Event-related design | 28 (*n.r.*) | English | 15 (2) | 10−17 | right | I > C | Yes | 102 |
| 20 | Schulte | 2020 | Journal | Stroop task | Block design | 178 (87) | English | 16 (2.3) | 12−21 | *n.r.* | I > C | *n.r.* | 10.35 |
| 21 | Sebastian | 2021 | Journal | emotional Simon task | Block design | 58 (*n.r.*) | English | 14 (1.68) | 10−16 | *n.r.* | I > C | Yes | 59 |
| 22 | Sheridan | 2014 | Journal | Simon task | Block design | 33 (19) | English | 8.1 (1.66) | 5.7−10.7 | *n.r.* | I > C | *n.r.* | 44 |
| 23 | Tamm | 2002 | Journal | Stroop task | Event-related design | 14 (*n.r.*) | English | 15.43 (3.79) | 10−22 | *n.r.* | I > N | Yes | 76.16 |
| 24 | Vaidya | 2005 | Journal | Flanker task | Event-related design | 10 (7) | English | 9.2 (1.3) | 7-11 | *n.r.* | I > N | No | 43.3 |
| 25 | van't Ent | 2009 | Journal | Stroop task | Event-related design | 18 (*n.r.*) | English | 12 (*n.r.*) | 7-12 | *n.r.* | I > C | Yes | 61.7 |
| 26 | Wang | 2009 | Journal | Stroop task | Event-related design | group1: 22 (15) | English | 15 (1.1) | 13−17 | *n.r.* | I > C | *n.r.* | 29.7 |
| 27 | Wang | 2009 | Journal | Stroop task | Event-related design | group2: 22 (18) | English | 14.8 (1.2) | 13−17 | *n.r.* | I > C | *n.r.* | 31.4 |
| **Young to middle-aged adults** | | | | | | | | | | | | | |
| 1 | Aarts | 2008 | Journal | Simon task | Event-related design | 12 (2) | English | 21.2 (*n.r.*) | 18−24 | right | I > C | Yes | 94.85 |
| 2 | Adleman | 2002 | Journal | Stroop task | Event-related design | 11 (3) | English | 19.98 (1.72) | 17.39−22.68 | right | I > C | *n.r.* | *n.r.* |
| 3 | Ansari | 2006 | Journal | number-size congruity task (Stroop-like) | Event-related design | 14 (6) | English | 21 (*n.r.*) | 18−24 | right | I > C | *n.r.* | 38.4 |
| 4 | Balodis | 2013 | Journal | Stroop task | Event-related design | 35 (16) | English | 38.4 (*n.r.*) | 19−64 | *n.r.* | I > C | *n.r.* | 218.46 |
| 5 | Barros-Loscertales | 2011 | Journal | Stroop task | Block design | 16 (*n.r.*) | English | 34.2 (8.86) | *n.r.* | right | I > N | Yes | 55.12 |
| 6 | Basten | 2011 | Journal | Stroop task | Event-related design | 46 (23) | English | 22.3 (2) | 19−27 | right | I > C | *n.r.* | 48.67 |
| 7 | Brass | 2005 | Journal | Stroop task | Event-related design | 20 (8) | English | 26 (*n.r.*) | 21−37 | right | I > C | *n.r.* | 163.1 |
| 8 | Bunge | 2002 | Journal | Flanker task | Event-related design | 10 (5) | English | 27 (*n.r.*) | 18−44 | right | I > N | Yes | 23 |
| 9 | Bush | 2003 | Journal | Simon task | Block design | 8 (*n.r.*) | English | 30.4 (5.6) | *n.r.* | right | I > C | *n.r.* | 308 |
| 10 | Bush | 1998 | Journal | Stroop task | Block design | 9 (5) | English | 24.2 (2.3) | *n.r.* | right | I > N | *n.r.* | 46 |
| 11 | Carp | 2012 | Journal | Simon task | Event-related design | 21 (*15*) | English | 39.8 (*n.r.*) | 23−51 | right | I > C | Yes | 237.3 |
| 12 | Carter | 1995 | Journal | Stroop task | Event-related design | 15 (*n.r.*) | English | 34.3 (*n.r.*) | 22−49 | right | I > N | Yes | 97 |
| 13 | Christensen | 2011 | Journal | Stroop task | Event-related design | 26 (10) | English | 25.9 (*n.r.*) | 19−53 | right | I > N | Yes | 38.5 |
| 14 | Cieslik | 2010 | Journal | Simon task | Event-related design | 24 (13) | English | 29 (*n.r.*) | 20−59 | right | I > C | Yes | 62.75 |

| | | | | | | | | | | | | | |
|---|---|---|---|---|---|---|---|---|---|---|---|---|---|
| 15 | Coderre | 2008 | Journal | Stroop task | Block design | 9 (2) | English | 36 (9.4) | *n.r.* | right | I > C | *n.r.* | 94.7 |
| 16 | DeVito | 2012 | Journal | Stroop task | Event-related design | 12 (5) | English | 31.0 (8.6) | 18−50 | right | I > C | Yes | 1323 |
| 17 | Durston | 2003 | Journal | Flanker task | Event-related design | 9 (5) | English | 25.7 (*n.r.*) | *n.r.* | right | I > C | Yes | 64 |
| 18 | Fan | 2003 | Journal | Flanker task | Event-related design | 12 (6) | English | 24.7 (4.6) | 18−34 | right | I > C | Yes | 140 |
| 19 | Fan | 2008 | Journal | Flanker task | Event-related design | 16 (8) | English | 27.2 (5.7) | 18−36 | right | I > C | Yes | 101 |
| 20 | Fan | 2007 | Journal | Flanker task | Event-related design | 19 (10) | English | 26 (*n.r.*) | 18−59 | *n.r.* | I > C | Yes | 55 |
| 21 | Fechir | 2010 | Journal | Stroop task | Block design | 16 (*n.r.*) | English | 23.8 (1.4) | 21−26 | right | I > C | Yes | 70.1 |
| 22 | Forstmann | 2008 | Journal | Simon task | Block design | 24 (9) | English | 24.2 (2.76) | *n.r.* | right | I > N | *n.r.* | 21 |
| 23 | Fruhholz | 2011 | Journal | Simon & Flanker task | Block design | 24 (3) | English | 23.91 (5.31) | 18−43 | right | I > C | Yes | 58 |
| 24 | George | 1994 | Journal | Stroop task | Event-related design | 21 (11) | English | 38.4 (13.2) | *n.r.* | right | I > N | *n.r.* | 192 |
| 25 | Georgiou-Karistianis | 2012 | Journal | Simon task | Event-related design | 13 (9) | English | 33.7 (7.9) | *n.r.* | right | I > C | *n.r.* | 58 |
| 26 | Grandjean | 2013 | Journal | Stroop task | Block design | 25 (12) | English | 21.8 (2.68) | 18−29 | right | I > C | Yes | 182.05 |
| 27 | Harrison | 2005 | Journal | Stroop task | Hybrid block/Event-related design | 9 (7) | English | 27.4 (9.1) | *n.r.* | right | I > C | Yes | -181.2 |
| 28 | Hazeltine | 2003 | Journal | Flanker task | Event-related design | 10 (5) | English | 27 (*n.r.*) | 18−44 | right | I > N | Yes | 46 |
| 29 | Hazeltine | 2000 | Journal | Flanker task | Hybrid block/Event-related design | 8 (3) | English | 21 (*n.r.*) | 18−24 | right | I > C | *n.r.* | 39 |
| 30 | Ivanov | 2012 | Journal | Flanker task | Event-related design | 16 (10) | English | 30.63 (7.44) | 21−45 | right | I > C | Yes | 61.4 |
| 31 | Jiang | 2014 | Journal | Simon task | Block design | 21 (10) | English | 21.3 (*n.r.*) | *n.r.* | *n.r.* | I > C | Yes | 33 |
| 32 | Kerns | 2006 | Journal | Simon task | Block design | 26 (12) | English | 24.2 (4.5) | 18−36 | right | I > C | Yes | 16.8 |
| 33 | Kerns | 2005 | Journal | Stroop task | Block design | 13 (8) | English | 35.6 (8.9) | *n.r.* | right | I > C | Yes | 90.4 |
| 34 | Kim | 2012 | Journal | Simon task | Event-related design | 16 (7) | English | 23.6 (2.9) | 18−35 | right | I > C | Yes | -87 |
| 35 | Kim | 2014 | Journal | Stroop task | Event-related design | 18 (10) | English | 25.3 (3.6) | 19−34 | right | I > C | Yes | 205.8 |
| 36 | King | 2012 | Journal | Flanker task | Event-related design | 25 (11) | English | 23.8 (*n.r.*) | 18−33 | right | I > C | Yes | 109 |
| 37 | Korsch | 2014 | Journal | Simon task | Block design | 20 (10) | English | 22.95 (2.72) | *n.r.* | right | I > C | Yes | 37 |
| 38 | Kozasa | 2012 | Journal | Stroop task | Block design | 19 (9) | English | 43.8 (9.35) | *n.r.* | right | I > C | Yes | 87.8 |
| 39 | Krebs | 2015 | Journal | Stroop task | Event-related design | 14 (8) | English | 22.5 (*n.r.*) | 18−35 | both | I > (C+N)/2 | Yes | 20.8 |
| 40 | Laeng | 2011 | Journal | Stroop task | Hybrid block/Event-related design | 10 (0) | English | 53.8 (14) | *n.r.* | *n.r.* | I > C | *n.r.* | 61.8 |
| 41 | Li | 2015 | Journal | Simon task | Block design | 24 (13) | English | 23 (3.26) | *n.r.* | right | I > C | Yes | 117 |
| 42 | Lutcke | 2009 | Journal | Flanker task | Event-related design | 12 (3) | English | 28 (6) | *n.r.* | right | I > C | Yes | 66 |
| 43 | Mathis | 2009 | Journal | Stroop task | Block design | 12 (7) | English | 26.8 (3.4) | 22−30 | *n.r.* | I > N & I > C | Yes | 60 |
| 44 | Mathis | 2009 | Journal | Stroop task | Block design | 12 (4) | English | 51.7 (3.1) | 46–55 | *n.r.* | I > N & I > C | Yes | 63.5 |
| 45 | Matthews | 2004 | Journal | Stroop task | Block design | 18 (11) | English | 39 (*n.r.*) | 27−56 | *n.r.* | I > C | *n.r.* | 112 |
| 46 | McNab | 2008 | Journal | Flanker task | Event-related design | 14 (4) | English | 24 (*n.r.*) | 22−34 | right | I > C | *n.r.* | 30 |
| 47 | Mead | 2002 | Journal | Stroop task | Block design | 18 (8) | English | 26.7 (*n.r.*) | 18−46 | right | I > C | *n.r.* | 56 |
| 48 | Milham | 2002 | Journal | Stroop task | Block design | 12 (7) | English | 23 (*n.r.*) | 21−27 | right | I > C & I > N | *n.r.* | 147 |
| 49 | Mitchell | 2005 | Journal | Stroop task | Block design | 15 (4) | English | 23.3 (6.31) | *n.r.* | right | I > N | *n.r.* | 68 |
| 50 | Mitchell | 2010 | Journal | Stroop task | Block design | 28 (3) | English | 20.2 (2.9) | *n.r.* | right | I > C | *n.r.* | 82.5 |
| 51 | Nakao | 2005 | Journal | Stroop task | Event-related design | 14 (5) | English | 30.2 (5.13) | 24−43 | both | I > C | *n.r.* | 116 |
| 52 | Ochsner | 2009 | Journal | Flanker task | Event-related design | 16 (7) | English | 21.22 (*n.r.*) | *n.r.* | right | I > C | Yes | 25.43 |
| 53 | Page | 2009 | Journal | Simon task | Event-related design | 11 (*n.r.*) | English | 34.1 (10.1) | *n.r.* | right | I > C | Yes | 120.1 |
| 54 | Piai | 2013 | Journal | Stroop task | Event-related design | 23 (11) | English | 21.2 (*n.r.*) | 18−29 | right | I > C & I > N | Yes | 25 |
| 55 | Polosan | 2011 | Journal | Stroop task | Block design | 14 (4) | English | 35.9 (7.2) | *n.r.* | right | I > C | Yes | 113.23 |
| 56 | Pompei | 2011 | Journal | Stroop task | Block design | 48 (25) | English | 36.33 (12.8) | *n.r.* | *n.r.* | I > N | Yes | 250 |
| 57 | Rahm | 2014 | Journal | Stroop task | Block design | 11 (8) | English | 34.9 (7.8) | *n.r.* | right | I > N | *n.r.* | *n.r.* |
| 58 | Ravnkilde | 2002 | Journal | Stroop task | Event-related design | 46 (16) | English | 41 (11.6) | 21−65 | both | I > C | *n.r.* | 292.3 |
| 59 | Roberts | 2008 | Journal | Stroop task | Block design | 16 (9) | English | 24.3 (*n.r.*) | 16−42 | right | I > N | *n.r.* | 62 |
| 60 | Robertson | 2015 | Journal | number-size congruity task (Stroop-like) | Event-related design | 16 (8) | English | 23 (*n.r.*) | 19−27 | right | I > C | *n.r.* | 52.035 |
| 61 | Roelofs | 2006 | Journal | arrow word Stroop task | Event-related design | 12 (4) | English | 23 (*n.r.*) | 21−28 | right | I > C | Yes | 44.05 |
| 62 | Rubia | 2006 | Journal | Simon task | Block design | 23 (23) | English | 28 (6) | 20−43 | right | I > C | Yes | 127 |
| 63 | Schmidt | 2012 | Journal | Stroop task | Event-related design | 31 (14) | English | 24.125 (*n.r.*) | 22−32 | both | I > C | Yes | 148.25 |
| 64 | Schulze | 2013 | Journal | auditory Stroop task | Block design | 8 (3) | English | 24.8 (2) | 22−27 | right | I > C | Yes | *n.r.* |
| 65 | Sebastian | 2013 | Journal | Simon task | Event-related design | 49 (19) | English | 39.96 (17.14) | 20−77 | right | I > C | Yes | 55.62 |
| 66 | Sebastian | 2012 | Journal | Simon task | Event-related design | 24 (11) | English | 30.3 (8.1) | *n.r.* | right | I > C | Yes | 60.18 |
| 67 | Sebastian | 2013 | Journal | Simon task | Event-related design | 21 (12) | English | 24.24 (2.3) | *n.r.* | right | I > C | Yes | 58.19 |
| 68 | Sheu | 2012 | Journal | Stroop & MSIT task | Block design | 26 (14) | English | 40 (6) | *n.r.* | *n.r.* | I > C | *n.r.* | *n.r.* |

| | | | | | | | | | | | | | |
|---|---|---|---|---|---|---|---|---|---|---|---|---|---|
| 69 | Soeda | 2005 | Journal | Stroop task | Block design | 11 (7) | English | 28.1 (4.7) | 23−35 | right | I > C | *n.r.* | *n.r.* |
| 70 | Sommer | 2008 | Journal | Simon task | Block design | 12 (12) | English | 29.1 (*n.r.*) | 22−37 | right | I > C | Yes | 86.5 |
| 71 | Terry | 2012 | Journal | Stroop task | Event-related design | 20 (*n.r.*) | English | 20.4 (1.6) | 18−25 | right | I > C | *n.r.* | 76.81 |
| 72 | Ullsperger | 2001 | Journal | Flanker task | Event-related design | 12 (5) | English | 24.9 (*n.r.*) | 21−29 | right | I > C | Yes | 61 |
| 73 | Verstynen | 2014 | Journal | Stroop task | Event-related design | 30 (20) | English | 31 (*n.r.*) | 21−45 | both | I > N | Yes | 61.4 |
| 74 | Weiss | 2007 | Journal | Stroop task | Block design | 8 (*n.r.*) | English | 26.89 (3.1) | *n.r.* | right | I > C | *n.r.* | *n.r.* |
| 75 | Wittfoth | 2006 | Journal | Simon task | Event-related design | 20 (3) | English | 25.5 (*n.r.*) | 21−31 | *n.r.* | I > C | Yes | 50 |
| 76 | Wittfoth | 2008 | Journal | Simon task | Event-related design | 20 (3) | English | 25.5 (*n.r.*) | 21−31 | right | I > C | Yes | 56 |
| 77 | Ye | 2009 | Journal | Stroop task | Event-related design | 19 (7) | English | 21 (*n.r.*) | 19−23 | right | I > C | Yes | 85 |
| 78 | Zhu | 2010 | Journal | Flanker task | Event-related design | 22 (11) | English | 20 (3) | *n.r.* | *n.r.* | I > C | Yes | 130 |
| 79 | Zoccatelli | 2010 | Journal | Stroop task | Block design | 10 (8) | English | 28 (*n.r.*) | 22−40 | right | I > C | Yes | 117.4 |
| 80 | Zurawska | 2011 | Journal | Flanker task | Event-related design | 18 (8) | English | 25.3 (*n.r.*) | 20−34 | right | I > C | Yes | 45 |
| 81 | Zysset | 2007 | Journal | Stroop task | Event-related design | 47 (23) | English | 42 (*n.r.*) | 22−75 | *n.r.* | I > N | Yes | 90.3 |
| 82 | Chen | 2022 | Thesis | Stroop task | Event-related design | 41 (19) | Chinese | 21.29 (2.83) | *n.r.* | right | I > C | Yes | 67 |
| 83 | Cui | 2011 | Thesis | Stroop task | Event-related design | group1: 10 (1) | Chinese | 21 (1.9) | 18−28 | right | I > C | *n.r.* | 136 |
| 84 | Cui | 2011 | Thesis | Stroop task | Event-related design | group2: 10 (10) | Chinese | 22.4 (3.0) | 18−28 | right | I > C | *n.r.* | 143 |
| 85 | Mou | 2018 | Thesis | Stroop task | Event-related design | 37 (16) | Chinese | 21.4 (1.4) | *n.r.* | right | I > C | Yes | 32.5 |
| 86 | Qian | 2020 | Journal | Stroop task | Block design | 17 (8) | Chinese | 25 (2) | 22−28 | right | I > C | *n.r.* | 95.785 |
| **The elderly** | | | | | | | | | | | | | |
| 1 | Chuang | 2014 | Journal | Flanker task | Event-related design | 60 (15) | English | 64.6 (3.7) | 60−74 | right | I > C | Yes | 60.75 |
| 2 | Dash | 2019 | Journal | Flanker task | Event-related design | 18 (*n.r.*) | English | 73.94 (2.8) | *n.r.* | *n.r.* | I > C | Yes | 40.98 |
| 3 | Fernandez | 2019 | Journal | Flanker task | Event-related design | 34 (*n.r.*) | English | 72.7 (5.3) | *n.r.* | right | I > C | Yes | 148.6 |
| 4 | Gianaros | 2007 | Journal | Stroop task | Block design | 46 (*n.r.*) | English | 68.04 (1.35) | *n.r.* | *n.r.* | I > C | *n.r.* | 18.07 |
| 5 | Gordon | 2015 | Journal | Stroop task | Event-related design | 71 (37) | English | 63.5 (*n.r.*) | 49−78 | right | I > C | *n.r.* | 94 |
| 6 | Huang | 2012 | Journal | physical Stroop task | Event-related design | 18 (9) | English | 66.1 (4.15) | 61−73 | right | I > C | Yes | 107.5 |
| 7 | Korsch | 2014 | Journal | Flanker & Simon task | Event-related design | 19 (10) | English | 70.26 (3.49) | *n.r.* | right | I > C | Yes | 35 |
| 8 | Mathis | 2009 | Journal | Stroop task | Block design | 12 (9) | English | 62.8 (3) | 60−68 | *n.r.* | I > N & I > C | Yes | 81 |
| 9 | Milham | 2002 | Journal | Stroop task | Block design | 10 (7) | English | 68 (*n.r.*) | 60−75 | right | I > C | *n.r.* | 220 |
| 10 | Nagamatsu | 2011 | Journal | Flanker task | Event-related design | 73 (0) | English | 69.6 (3.1) | 65−75 | *n.r.* | I > C | Yes | *n.r.* |
| 11 | Onur | 2011 | Journal | visual-spatial Stroop & Simon-like task | Event-related design | 13 (8) | English | 63.81 (6) | *n.r.* | right | I > C | *n.r.* | 48.2 |
| 12 | Prakash | 2009 | Journal | Stroop task | Event-related design | 25 (*n.r.*) | English | 65.5 (*n.r.*) | 58−75 | right | I > N | Yes | 89.18 |
| 13 | Puente | 2014 | Journal | Stroop task | Event-related design | 26 (10) | English | 74 (5.5) | 65−85 | both | I > C | Yes | 240 |
| 14 | Rizio | 2017 | Journal | Stroop task | Event-related design | 20 (*n.r.*) | English | 67 (*n.r.*) | 60−79 | right | I > N | Yes | 80.77 |
| 15 | Won | 2019 | Journal | Flanker task | Event-related design | 32 (8) | English | 66.2 (7.3) | 55−80 | right | I > C | *n.r.* | 85.2 |
| 16 | Zhu | 2010 | Journal | Flanker task | Event-related design | 22 (9) | English | 74 (6) | *n.r.* | *n.r.* | I > C | Yes | 206 |

Note. *n.r.* = not reported; I = incongruent condition; C = congruent condition; N = neutral condition.
